# Robust conformational space exploration of cyclic peptides by combining different MD protocols and force fields

**DOI:** 10.1101/2025.03.31.646350

**Authors:** Samuel Murail, Jaysen Sawmynaden, Akli Zemirli, Maud Jusot, Fabio Pietrucci, Jacques Chomilier, Pierre Tufféry, Dirk Stratmann

## Abstract

Cyclic peptides are an important class of pharmaceutical drugs. We used here replica-exchange molecular dynamics (REMD) and simulated tempering (ST) simulations to explore the conformational landscape of a set of nine cyclic peptides. The N-ter to C-ter backbone cyclized peptides of seven to ten residues were previously designed for a high conformational stability with a mixture of L and D amino acids. Their experimental NMR structures were available in the protein data bank (PDB). For each peptide we tested several force fields, Amber96, Amber14, RSFF2C and Charmm36m in implicit and explicit solvent. We find that the variability of the free energy maps obtained from several protocols is larger than the variability obtained by just repeating the same protocol. Running multiple protocols is therefore important for the convergence assessment of REMD or ST simulations. The majority of the free energy maps showed clusters with a high RMSD to the native structures, revealing the residual flexibility of this set of cyclic peptides. The high RMSD clusters had in some cases the lowest free energy, rendering the prediction of the native structure more difficult with a single protocol.

Fortunately, the combination of four implicit solvent REMD and ST simulations, mixing the Amber96 and Amber14 force fields, predicted robustly the native structure.

As implicit solvent in the REMD or ST setup are up to one hundred times faster than explicit solvent simulations, so running four implicit solvent simulations is a good practical choice. We checked that the use of an explicit solvent REMD or ST simulation, taken alone or combined with implicit solvent simulations, did not significantly improved our results. It results that our combination of four implicit solvent simulations is tied in terms of success rate with much more expensive combinations that include explicit solvent simulations. This may be used as guideline for further studies of cyclic peptide conformations.

## Introduction

Peptides have attracted increasing attention over the past decade as a viable alternative to small molecules for developing drug molecules capable of interfering with protein-protein interactions particularly well. ^3–7^ While linear peptides composed of only natural amino acids suffer from degradation, cyclic peptides display a stronger resistance to the proteases. ^8^ In addition, their conformational space is usually more restricted than that of linear peptides, ^9^ reducing the entropy loss upon binding, and leading to a stronger binding affinity. ^10,11^ Cyclisation is usually achieved by a peptide bond between the N-ter and the C-ter amino acid or by disulfide bonds between two cysteine residues. Combining both approaches allows the design of hyperstable cyclic peptides. ^12^ Conformational stability can also be obtained by a combination of L and D-amino acids, ^13^ without the need for disulfide bonds at least for small cyclic peptides up to about ten residues. ^2^ Combining the cyclisation with chemical modifications, like non-canonical amino acids ^14,15^ or N-methylation, can increase the specificity to the target ^16^ and reduce the conformation flexibility. ^17^ Cyclic peptides that incorporate chemical modifications are named *macrocycles*, they represent a class of drugs on their own right, ^18^ with an important field of research. The chemical space available to macrocycles is huge and as for small molecules, virtual libraries of billions of compounds have been developed. ^19^ To interfere with intracellular targets, drugs must pass the cell membrane. Fortunately, cyclic peptides can pass this barrier, as documented in the recent database CycPeptM-PDB gathering experimental membrane permeability measurements of over 7000 cyclic peptides, of which more than the half present a high membrane permeability, ^20^ and *in-vitro* workflows have been developed recently for generating orally available cyclic peptides. ^21^ Overall, there is a bright future for cyclic peptides and macrocycles in therapeutic drug development ^22^ with an average of about one cyclic peptide drug approved per year. ^7^ The numerous possibilities for in-vitro and in-silico design approaches for peptide therapeutics in general are summarized in several recent reviews. ^23–29^

While cyclic peptides are less flexible than linear peptides, some degree of flexibility remains depending on the length and the composition of the amino acid sequence, as well as the use of chemical modifications. ^17,30^ In a drug design context, it is important to be able to access the entire conformational landscape accessible to a cyclic peptide, for two main reasons: First, a poorly flexible cyclic peptide will likely lose less conformational entropy upon binding than a more flexible peptide. ^10,11^ To quantify this, the accessible conformational space can be a possible indicator. Second, a flexible cyclic peptide has usually a different conformation in free form than in its bound form inside the protein-peptide complex, due to the additional protein-peptide interactions. ^31^ For an optimal binding affinity, it is important that the bound form is already explored inside the conformational ensemble of the free form (*conformational selection*) or that the free form can be converted into the bound conformation (*induced fit*) without having to cross a too high free energy barrier. ^32^ For some applications, a high flexibility is needed. For example, it is sometimes necessary to travel “‘a tunnel”‘ to the binding site. So cyclic peptides cannot always be designed as “‘rocks”‘, but have to incorporate an appropriate level of flexibility. Experimental structure determination methods like NMR or X-ray crystallography will resolve the lowest free energy structure, and perhaps a second minor conformation, but not the whole conformational space and the free energy barriers among several meta-stable conformations. Therefore conformational space explorations are usually done with in-silico methods.

The conformational space explored by cyclic peptides can be characterized *in-silico* with molecular dynamics (reviewed in ^33^), Monte-Carlo simulations and other sampling methods. The Rosetta software suite ^34^ uses Monte-Carlo simulations and kinematic closure algorithms to sample cyclic peptides conformations. ^12^ A similar kinematic closure algorithm has been implemented in EGSCyP (Exhaustive Grid Search for Cyclic Peptides) ^35^ that uses an exhaustive exploration for cyclic pentapeptides composed of standard, D-amino acid and N-methylated amino acids. Conventional molecular dynamics simulations (cMD) are in general not sufficient to pass the high free energy barriers that separate different conformations of cyclic peptides. Therefore, enhanced sampling molecular dynamics simulations are used, like replica-exchange MD, ^35–43^ bias-exchanged metadynamics (BE-MetaD). ^41,42,44–53^ Machine learning can even accelerate further the exploration of the conformational space of cyclic peptides. ^54,55^ AlphaFold2^56^ has also been applied to cyclic peptides, ^57–59^ but it is limited by its design to the prediction of the 3D structure and not to the identification of a conformational ensemble. Whereas AlphaFold2 is limited to natural amino acids, the recent AlphaFold3^60^ may be used to predict the structure of cyclic peptides that incorporate modified amino acids, as it can be applied to any small molecule in general, but this remains the subject for furhter investigations.

In the present study, we assess the performance and convergence of several force fields in molecular dynamics (MD) simulations for the exploration of the conformational space of a set of nine cyclic peptides, ranging from seven to ten amino acids. We run in parallel two enhanced sampling methods, replica exchange molecular dynamics (REMD) and simulated tempering (ST), to exclude bias from either sampling method.

A very recent study ^61^ of the Yu-Shan Lin group tested several force fields on a set of 12 smaller cyclic peptides, ranging from five to seven amino acids. The smaller size stabilized the eight sequences without D-amino acids. The remaining four pentapeptides had one or two D-amino acids. They performed BE-MetaD simulations in explicit solvent for most of the force fields tested. In our study we focused on implicit solvent simulations to assess if this kind of solvent model is sufficient for the elucidation of the conformational space of free-form cyclic peptides. We focused also on medium sized cyclic peptides, as the conformational space can be more difficult to be resolved. Therefore our study is complementary to the similar study ^61^ of the Yu-Shan Lin group.

## Materials & Methods

### Cyclic peptide dataset

Most cyclic peptides present in the Protein Data Bank (PDB) are bound with other molecules or have chemical modification (like N-methyl). In the present study we choose a set of nine cyclic peptides available in free form in the PDB and having a medium size from seven to ten amino acids. All but one are peptides with a mixture of L and D amino acids, optimized for conformational stability by the David Baker group using Rosetta. ^2^ We employ here the same peptide names as in their paper, like “‘7.B”‘ for a peptide that has seven amino acids and that is the second design of this size (see table 1). The peptide 8.C in table 1 is not present in the study of Hosseinzadeh et al. It is composed of standard amino acids only. ^1^ We added it to see if a cyclic peptide of this size can be as stable as the peptides that are stabilized by D-amino acids and prolines.

**Table 1:**
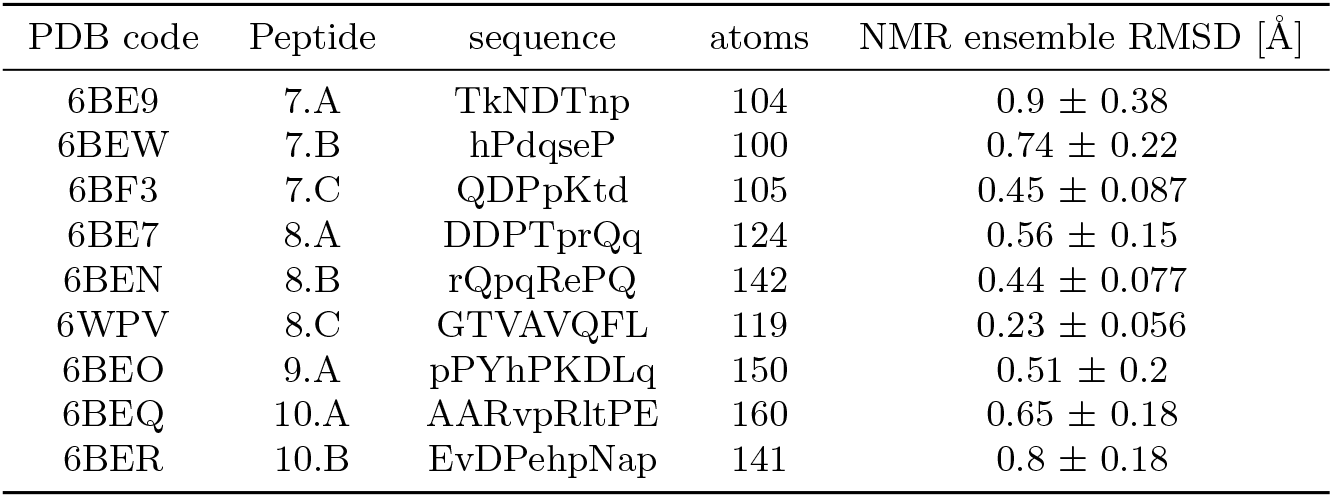
Cyclic peptide data set. All structures of the peptides in free form (i.e. not bound to a protein) have been solved by NMR.. Upper case letters are L-amino acids and lower case letters are D-amino acids. The all L-amino acids peptide 8C was published in Ref., ^1^ whereas the other peptides were published in Ref. ^2^ As reference structure we used the best representative conformer of the NMR ensemble which is here the first model for all nine PDB files. The conformational variability among the NMR ensemble is quantified with the average RMSD of the 19 models compared to the first model.

### Implicit solvent REMD protocol

Our REMD protocol is similar as the one we used in a former study on cyclic pentapeptides ^35^ and inspired from the protocol presented in the work of Wakefield et al. ^40^

The starting structures of the cyclic peptides were constructed manually with ChimeraX. ^62^ Topology files were generated with Ambertools ^63^ and converted in GROMACS format with acpype. ^64^

Temperature REMD simulations were setup with eight replica with OBC (Onufriev, Bashford, and Case) ^65^ GBSA implicit solvent and run with GROMACS 5.1.2, ^66^ more recent GROMACS versions do not include implicit solvents any more. A short minimization was first done with an alternation of one step of steepest descent every 500 conjugate gradient steps. The maximum number of steps was set to 50,000 with a step size of 0.01 nm and an energy convergence criterion of 10 kJ/(mol*nm). The thermalizations and simulations were done in a temperature range from 300K to 455K. The temperature values of the eight replica were chosen to keep the probabilities of accepting exchanges the same (300, 318, 337.97, 358.81, 380.85, 404.27, 429.12 and 455.50K). The acceptance ratio was high, as 55 to 65% of all exchange attempts were successful, thanks to the small number of atoms (100 to 150). Only a short phase of thermalization in NVT was realized during 100 ps (50,000 steps, 2 fs time step). The modified Berendsen thermostat was employed with the time constant *τ* = 0.1 ps.

A good sampling in REMD depends on the time step between exchange attempts between neighbouring replicas. ^67^ No consensus choices exist for this parameter. The time step used in the literature is between 0.01 to 100 ps ^36 68^ and it depends on the system and the process studied. Here we chose 1 ps as time step between exchange attempts. For each force field and peptide we produced trajectories of 1 *μ*s with a 2 fs time step for each replica, i.e. 8 *μ*s for each REMD simulation. The atomic coordinates were recorded every 10 ps. We repeated each REMD simulation five times (run 1 to 5) to access convergence. In total we produced in this study with the REMD protocol 8*μ*s * 5 runs * 9 peptides * 3 force fields = 1080 *μ*s of cumulative trajectory time (see table 2).

**Table 2:**
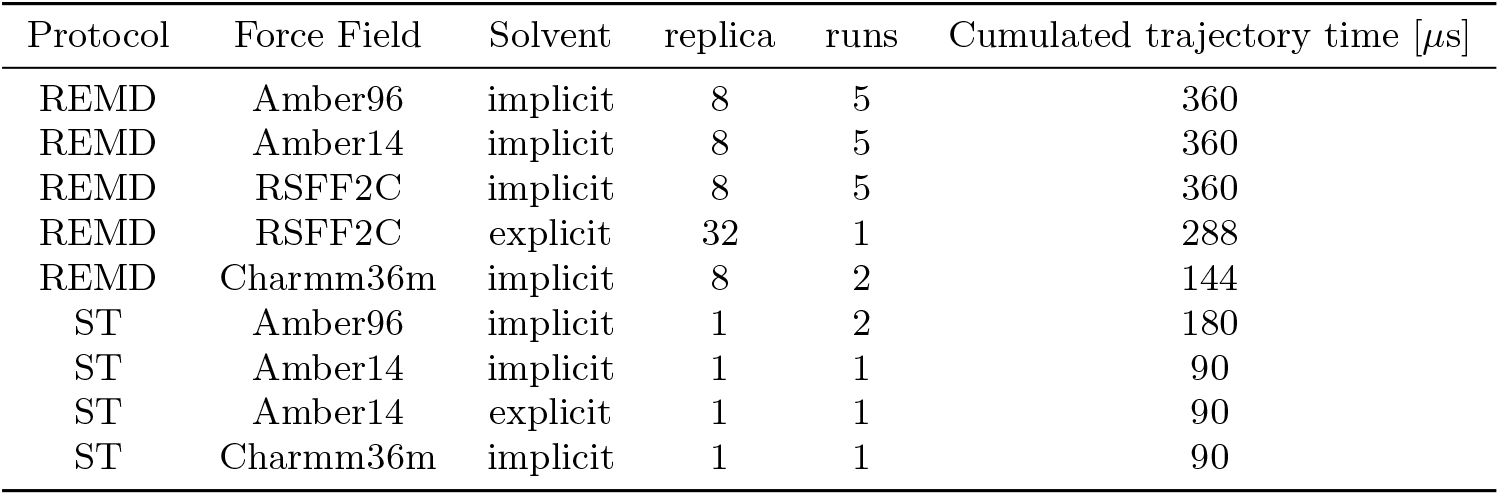
List of the nine combinations of sampling protocol, force field and solvent model that have been run in the present study. Each combination was run on the nine peptides from table 1. The trajectory length for REMD is 1*μ*s and for ST 10*μ*s. The cumulated trajectory time is the product of this trajectory length with the number of replica and with the number of repeated runs. The total cumulated trajectory time of the implicit and explicit solvent runs is 1584*μ*s and 378*μ*s, respectively.

| Protocol | Force Field | Solvent | replica | runs | Cumulated trajectory time [$\mu\text{s}$] |
| --- | --- | --- | --- | --- | --- |
| REMD | Amber96 | implicit | 8 | 5 | 360 |
| REMD | Amber14 | implicit | 8 | 5 | 360 |
| REMD | RSFF2C | implicit | 8 | 5 | 360 |
| REMD | RSFF2C | explicit | 32 | 1 | 288 |
| REMD | Charmm36m | implicit | 8 | 2 | 144 |
| ST | Amber96 | implicit | 1 | 2 | 180 |
| ST | Amber14 | implicit | 1 | 1 | 90 |
| ST | Amber14 | explicit | 1 | 1 | 90 |
| ST | Charmm36m | implicit | 1 | 1 | 90 |

To produce 1 *μs* per replica the cost is between 1000 to 2000 CPU-hours per REMD simulation. So in total we consumed about 5 runs * 9 peptides * 3 force fields * 1000-2000 CPU-hours = 135k-270k CPU-hours, which is quite reasonable thanks to the use of an implicit solvent.

Four force fields were tested with the REMD protocol: Charmm36m, ^69^ Amber96, ^70^ Amber14SB ^71^ and the coil-library-based Residue Specific Force Field RSFF2C ^72^ which is derived from the Amber14SB force field and has shown better performance for peptides, ^61^ proteins and loops. ^73^ We included the older Amber96 force field, as we employed it in our former study ^35^ and as it showed still a good performance on cyclic peptides.

### Explicit solvent REMD protocol

We setup also an explicit solvent REMD protocol with the RSFF2C force field, which has been tested only in explicit solvent in previous studies. Using the standard TIP3P explicit solvent model, a cubic water box with a minimum distance between the peptide and the box walls of 1.0 nm was equilibrated with GROMACS 2018.8. ^66^ In the case that the peptide has not a zero net electric charge, the total charge of the peptide + water system has been neutralized by adding sodium or chloride ions to the solvent. The energy of the whole system was then minimized with a steepest descent algorithm and each replica was equilibrated to its temperature in two steps, first with a simulation of 100 ps in the NVT ensemble followed by a simulation of 200 ps in the NPT ensemble. The time step of the equilibration and production simulations was set to 2 fs. A non-bonded cutoff of 1.0 nm was used with the Particle Mesh Ewald (PME) algorithm for long range electrostatic interactions. The modified Berendsen thermostat was employed with the time constant *τ* = 0.1 ps and the Parrinello-Rahman pressure coupling in the NPT ensemble with the time constant *τ* = 2.0 ps.

We employed 32 replica, as this value fitted well with our computer architecture. The temperature values were generated with the web-server ^74^ using the algorithm published in Ref. ^75^ using the same temperature range as for the implicit solvent protocol (see supplementary materials and methods for the temperatures list).

We obtained an acceptance ratio of 25 to 35% for the replica exchanges, depending on the number of atoms, which varied from 4300 to 6900 atoms (solvent + peptide) in function of the size of the peptide. Usually an acceptance ratio of around 30% is targeted and we were pleased to see that the final acceptance ratios were about twice the values (12-15%) predicted by the web-server mentioned above.

With two 32 cores CPUs per replica we obtained 50 to 75 ns/day on two recent 32 cores AMD EPYC 7452 @ 2.35GHz CPUs. Running all nine peptides on eighteen 32 cores CPUs in parallel, we obtained after about 13 to 20 days a single 1000 ns trajectory per peptide. In total this single run consumed about 200k CPU hours (15 days * 64 cores * 9 peptides). Repeating the simulation with five runs like for the implicit solvent was out of reach for this study, neither doing the whole benchmark of the three force fields in explicit solvent with REMD, as taken together it would have consumed 15 times more CPU hours, i.e. about 3 million CPU hours.

### ST simulations

Protein structure were prepared using the pdbfixer module of the OpenMM package, where a bond was specifically added in the peptide topology between the N backbone atom of the first residue and the C backbone atom of the last residue. ^76^ For simulation run with explicit solvent, peptides were solvated in truncated octahedron boxes with a padding of 1.5 nm and the TIP3P water model. Counter ions were added to counter the charge of the peptide, together with a 150 mM concentration of Na^+^Cl^-^ ions. Simulation were conducted with the OpenMM 7.7 Molecular Dynamics software. ^76^

Energy minimization was run for up to 10.000 steps, and followed by an equilibration step of 10 ns in the NPT ensemble. During production and equilibration, a 4 fs integration time step was used with the help of constraining all bonds involding a hydrogen atom, as well as using heavy hydrogens assigned to a 3 atomic mass units. A non-bonded cutoff of 1.0 nm was used with the Particle Mesh Ewald (PME) algorithm for long range electrostatic interactions. A pressure of 1.0 bar was applied during equilibration and production using the Monte-Carlo algorithm and a 25 time steps interval. Temperature was equilibrated using the Langevin Middle integrator and 1 ps^-1^ friction coefficient.

During the equilibration the temperature was fixed to 300.0 K. Simulated Tempering (ST) simulation were conducted using a protocol inspired from a previous work ^77^ using a python script written by Peter Eastman and modified to implement weight calculation of Park and Pande ^78^ and on-the-fly weight calculation of Nguyen et al. ^79^ During ST simulations, temperature exchanges were attempted every 10 ps and depending with the size of the peptide 15 to 20 temperature ladders spaced exponentially between 300.0 and 500.0 K were used. The ST simulation production run had a length of 10 μs.

Simulation run with implicit solvent were run in the NVT ensemble with no cutoff for non-bonded interaction. We used only 8 temperature ladders, and the temperature exchange were attempted every 6 ps. Implicit solvent models were GBN2^80^ for simulation run with Amber14 force field, as we used OBC model ^81^ with Amber96 and Charmm36m force field.

### Analysis of REMD and ST simulations

All analyses in this article were done on the REMD trajectory at 300K and the parts of the ST trajectory that were at 300K. We skipped the first 10% of the trajectory to reduce the bias of the starting conformation. We compared each frame to the experimental reference structure which is here the best representative conformer of the NMR ensemble (first model here). Two measures were calculated for each frame: the backbone RMSD to the experimental reference structure and the radius of gyration. Both were calculated with the mdtraj ^82^ python package on the heavy atoms of the backbone (N, *C*_*α*_, C and O). Free energy maps in function of both metrics were calculated and plotted with python using a custom function adapted from the PyEMMA package. ^83^ Therefore a 2D histogram is generated with numpy (100 bins for each axis) and translated into free energy values according the formula: *F* = −*kT* * log(*ρ*), with *ρ* being the density at each point of the free energy map.

### Convergence analysis

To determine the convergence of a MD simulation is not a trivial problem. As MD is a random walk, short simulations cannot sample enough conformational landscape and explore all free energy minima. Moreover the populations of the clusters can change during longer trajectories, as shown in figure 1a).

**Figure 1.**
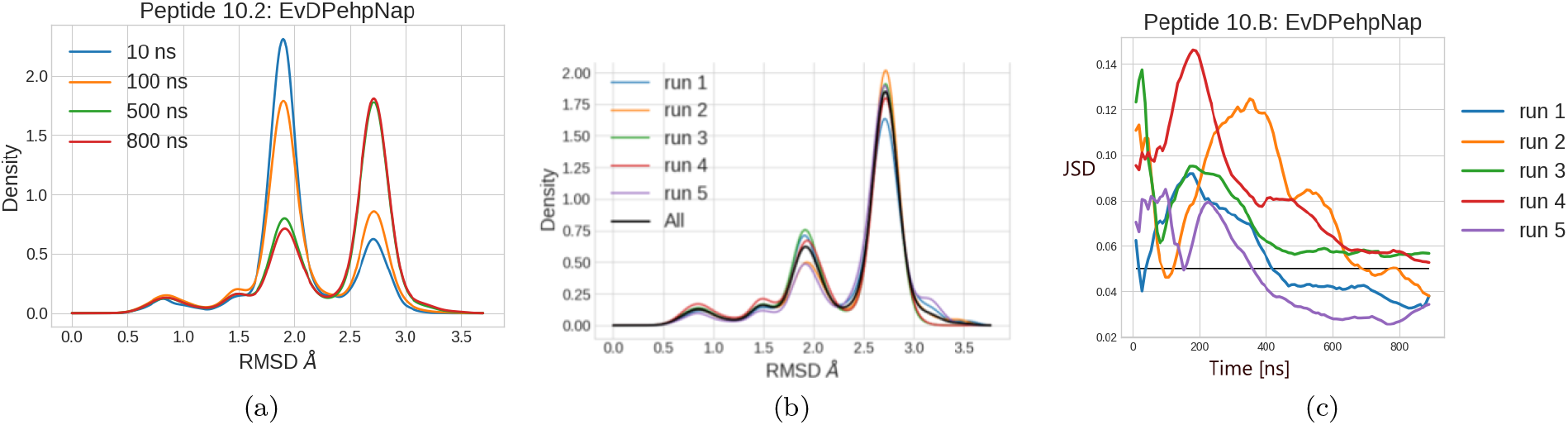
(a) REMD is a random walk, conformational space exploration is not the same with short (*<*100 ns) and long (*>* 500 ns) simulations. The determination of the simulated time necessary to get a converged simulation is important to conclude which clusters are most populated. In this example the population of the clusters converged after 500 ns. The RMSD values of the shown profiles are calculated to the experimental reference structure. (b) Comparison of the RMSD profiles of five REMD runs of 1*μs* (first 10% skipped) to the mean RMSD profile which combines all five profiles.(c) The Jensen-Shannon divergence (JSD) is used as metric to quantify the difference between two profiles. If the JSD value is above 0.05, we consider that the RMSD profile is different to the mean RMSD profile. Convergence of all five REMD runs is reached if all five curves are below 0.05, here only three of five runs converged, but the other two runs (run 3 and 4 here) are not far from the chosen threshold.

To assess quantitatively the convergence of our implicit solvent REMD simulations, we performed five independent runs for each peptide and force field combination. If the RMSD density profiles of all five runs superpose within a certain error margin, we consider that the REMD simulations converged. The RMSD values are calculated to the experimental reference structure, but in principle any structure could be used as reference here. To quantify this superposition we build first a mean density profile combining all five runs (see figure 1b)). To quantify how much each RMSD profile differ from the mean RMSD profile, the Jensen–Shannon divergence (JSD ^84^) is used:

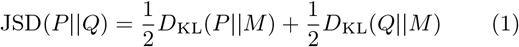

with *P* : RMSD density profile, *Q*: Mean RMSD density profile, 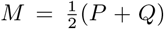 and *D*_KL_: Kullback-Leibler (KL) divergence.

Unlike KL, JSD is a symmetric metric (JSD(*A, B*) = JSD(*B, A*)) and the range of values is between 0 (similar profiles) to 1 (completely different profiles). If the JSD value is above 0.05, we consider that the RMSD profile is different to the mean RMSD profile (see figure 1b) as an example of JSD values around 0.05).

For each REMD run 100 RMSD density profiles are calculated along the trajectory (0-*t*_1_, 0-*t*_2_, 0-*t*_3_, …, 0-*t*_100_) to capture the simulation time evolution of the convergence, which allowed us to plot the JSD curves shown in figure 1c).

### 2D projections of the conformational landscape

The conformation landscape is projected here to a 2D map, using two complementary projections: the first one has on the x-axis the radius of gyration of the cyclic peptide backbone and on the y-axis the RMSD to the experimental reference structure calculated over the backbone atoms. This projection gives a good idea where the highest populated conformations are situated compared to the experimental reference structure. The radius of gyration is a good measure for the overall shape of the cyclic peptide ring. The second projection is obtained through a principal component analysis (PCA) of the cyclic peptide backbone atoms positions of the molecular dynamics trajectory (see below for more details). We employed the “‘MDanalysis”‘ python package ^85^ for the PCA calculation.

For the identification of low free energy clusters we employed two techniques: k-means clustering ^86^ and HDBscan clustering. ^87^ See supplementary materials and methods for more details.

### Prediction of the experimental reference structure

Although our molecular dynamics simulations yield an ensemble of conformations and not just a single structure, the most populated cluster of conformations is expected to be near to the experimental reference structure. This can be evaluted immediately by looking on the 2D maps that project the conformational landscape onto the RMSD to the experimental reference structure, by reporting the RMSD of the centroid structure of the most populated cluster, i.e. the one with the lowest free energy in general (not always, as the HDBscan clustering algorithm can merge together two clusters of low free energy that do not include the lowest free energy peak).

In real cases the experimental reference structure is in general not available. Therefore we setup a protocol to predict it without the use of the RMSD/radius of gyration projection, but by performing a principal component analysis (PCA) directly from the coordinates of the backbone atoms sampled during the MD simulation (REMD or ST protocol). To do a PCA, the MD trajectory needs first to be aligned to one reference frame. We simply choose the first frame as reference frame. From the PCA we retained as many components as necessary to get a cumulated variance of 80% of the total variance. In our case about four to six principal components (PC) had to be retained. We clustered this multidimensional PCA space with the HDBscan algorithm using the same parameters as described above. The PCA space is projected onto the first two principal components to give the 2D maps shown in figure S4. The centroid of the most populated and lowest free energy cluster is indicated by a green star. In the rare case that the position of the lowest free energy value in the PCA maps is different from the position of the most populated cluster, we plotted a cyan star for the former and a yellow star for the latter. For each star, its RMSD to the experimental reference structure and radius of gyration are calculated, and this coordinate is indicated again by a star of the same color on the 2D maps shown in figure 9.

We tested also a variant of this protocol, where the frame number corresponding to the centroid of the most populated cluster from the first PCA run is used as reference frame to realign the MD trajectory and to perform a second PCA run, that should be less dependent on the arbitrary choice of first frame as reference frame from the first run. As the results did not change significantly (results not shown), we retained the first version of our protocol.

### Combining different force fields and simulation protocols

As will be shown in the results section, each force field or simulation protocol yields a slightly different free energy map for the same peptide. Even though different force fields cannot be combined directly, their resulting free energy maps can be combined to obtain *consensus* free energy maps which should give a more complete view of the conformational space of a given peptide. We combined simulations having either a different force field or a different simulation protocol (REMD vs ST, implicit vs explicit solvent) or both. These combinations were done by concatenating the trajectories of the simulations. As the number of frames of each trajectory is not the same for each simulation, we determined its minimum value and subsampled the other trajectories with the same number of frames, which yielded equal weights of each simulation that contributed to the concatenated trajectory. As before, we constructed the free energy maps from the concatenated trajectories and applied the same protocol described above for the prediction of the experimental reference structure. All possible combinations over the nine simulations were done, by joining together two, three or more simulations.

The performance of each combination in the prediction of the experimental reference structure is quantified by a single value, the median RMSD of the first clusters among all nine peptides of our dataset. .

## Results & Discussion

### Convergence assessed by repeating simulations

Usually convergence of molecular dynamics simulations can be assessed by repeating the same simulation with different initial conditions (initial velocities for example). We assessed the convergence of REMD simulations as shown for the Amber96 force field for two peptides in figure 2. For the other peptides the results are shown in the supplementary figure S1. We compared the density profiles of each of the 5 runs of the RMSD to the experimental reference structure to the average density profile. For almost all peptides the density profiles converged at the end of the simulation (1000 ns, or more exactly 900 ns as we reject the first 10% to avoid bias from initial conditions), as shown in figure 2 for the peptide 8.C (see figure S1 for the other peptides). In terms of convergence, the worst case was peptide 10.A, but as can be seen from figure 2 the ranking of the three major clusters is not affected by the slow convergence, only the absolute values of the cluster sizes did not converge at the end of the simulation. The speed of convergence can be assessed quantitatively by tracing the evolution of the profile-profile distance measured by the Jensen-Shannon distance (JSD) (figure 3). For the 8C peptide all five simulations converged after only 300 ns, as all Jensen-Shannon distances are under the threshold that we defined (0.05) and that correspond to very high similarity of the density profiles. Overall the six smaller peptides 7A to 8C show a much faster convergence than the three larger peptides 9A to 10B, as demonstrated by the JSD curves in figure S2b and the first line of mean JSD values at the end of the trajectories in figure 4.

**Figure 2.**
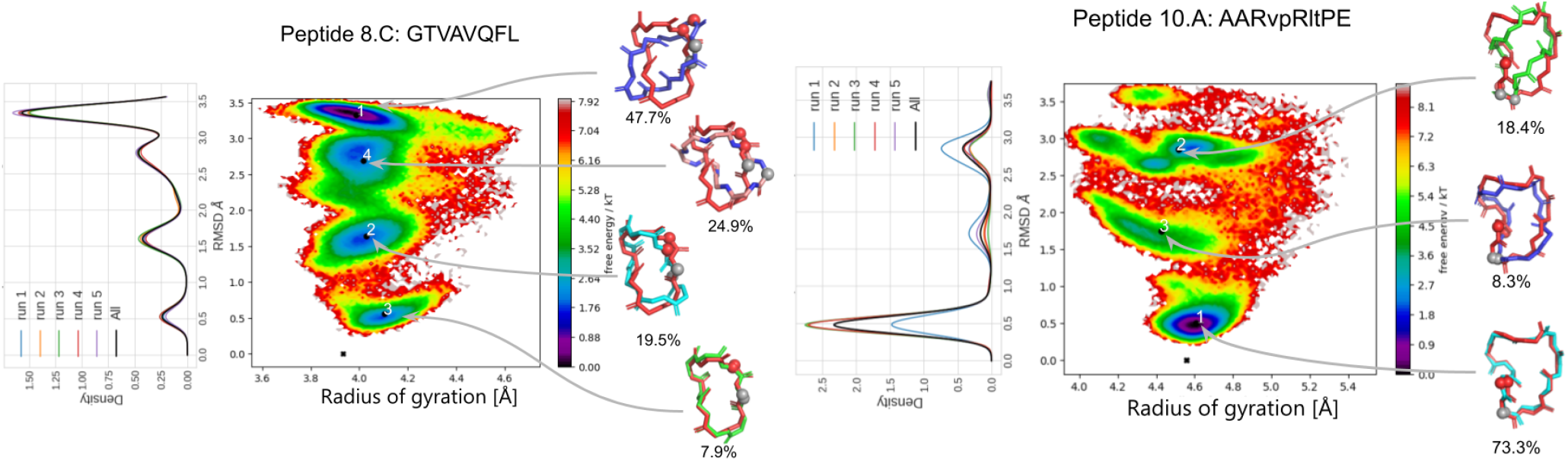
Convergence of simulations assessed by repeating simulations. The Amber96 force field in implicit solvent is employed here. Results for two peptides of REMD simulations repeated with five runs. For each run the density profile of the RMSD to the experimental reference structure is shown on the left. The density profile of the five concatenated trajectories is shown in black. The free energy map is also obtained by the assembly of all five runs. The map is clustered with the k-means algorithm and the centroid structures are superposed to the experimental reference structure (in red). The two first residues are indicated with a red and a gray ball. The size of each cluster is indicated below the 3D structures. For the other peptides the results are shown in the supplementary figure S1.

**Figure 3.**
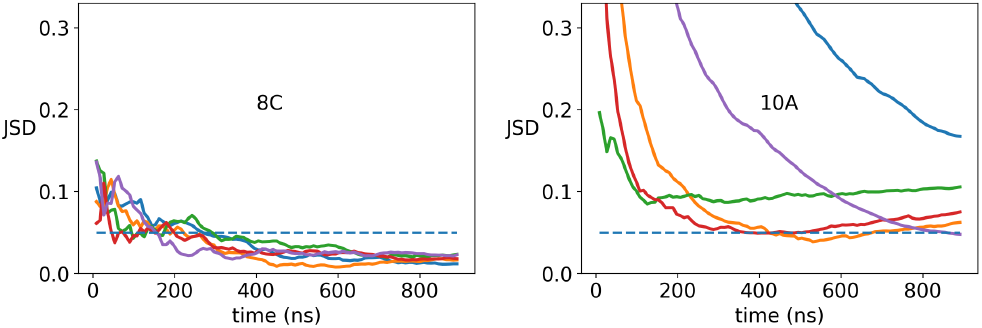
The speed of convergence is shown with the evolution of the Jensen-Shannon distance (JSD) of each RMSD density profile to the average RMSD density. Same REMD simulations as in figure 2. For the other peptides the time evolution of the JSD values is shown in figure S2b. Below the threshold of a JSD of 0.05 two RMSD profiles are nearly identical. Convergence is obtained, if all curves fall and stay below this threshold.

**Figure 4.**
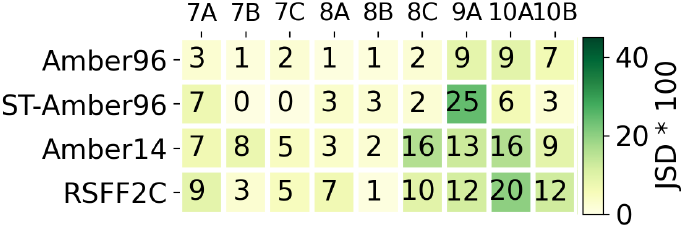
Individual convergence of REMD and ST simulations as measured by the mean JSD of all runs at the end of the trajectories (i.e. the mean of the last points of the curves in figure 3 and figure S2).

As the comparison of the 8C and 10A peptide results shows (figure 2), the convergence is not a guarantee to get the highest population for the cluster with the lowest RMSD to the experimental reference structure. Even though the REMD simulations of the 8C peptide converged very rapidly, all five runs yielded the highest population in a cluster more than 3Å away from the reference structure. In contrary, the five non converged runs of the 10A peptide all ranked the lowest RMSD cluster at the first position in terms of free energy/population size.

For the ST simulations we assessed the convergence by repeating the simulations with the Amber96 force field with two runs, run 1 and run 2. Figures 4 and S3 show that the convergence is here also very good among almost all peptides. Only one case showed a poor convergence (peptide 9A). This case is a bit surprising, as the cluster that accumulated 58% of the trajectory frames in run 1, did not even appear with a small percentage on the run 2. It seems that run 1 explored a cluster that has not been explored by run 2, whereas the other two clusters are both explored by each run. Among both runs we selected the run 1 for further analysis which is less favourable in terms of the ranking of the clusters, so the performance of ST with Amber96 is a bit lower rated for 9A. Unlike the REMD results, the convergence speed does not show a clear dependence on peptide size with the ST protocol. As can be seen in figure S2a the convergence speed is quite different among the nine peptides and also among the peptides with the same number of amino acids : peptide 7A shows a horizontal line above the 0.05 JSD threshold. 7B converged very rapidly and 7C needs about 4000 ns simulation time to converge both curves together. For the three larger peptides of eight amino acids, the convergence speed shows a similar heterogeneity. For the largest peptides of ten amino acids (10A and 10B), the convergence is average for 10B (JSD *<* 0.05 after 4500 ns) and slow for 10A due to a horizontal line like in the 7A case. Nevertheless a JSD threshold of 0.1 instead of 0.05 would classify both cases, 7A and 10A, as being converged after 6000 ns.

The individual convergence of REMD Amber96 and ST-Amber96 is summarized in figure 4 with the mean JSD values at the end of the trajectories. The convergence of the REMD Amber14 and RSFF2C simulations is shown in the same figure. The JSD values are higher here than for the Amber96 force field, especially for the larger or more flexible peptides (8C-10B), but still relatively contained. We cannot explain why the Amber14 and its derived RSFF2C force fields yield a lower convergence among repeated REMD simulations than the Amber96 force field. Maybe the fact that Amber14 and RSFF2C were developed for an explicit solvent plays a role here.

### Agreement across different simulation conditions

Even though each simulation protocol taken alone shows overall very good convergence properties, the agreement among different simulation protocols or force fields is not as good. A quick comparison of the results shown in figures 2 and S3 for peptide 8C illustrates this fact: while the most populated cluster of the REMD simulations is at about 3.3Å from the experimental reference structure, the ST simulations yield a better ranking with the most populated cluster at about 1.7Å. This is more due to different population sizes of each cluster than to “‘shape”‘ differences of the free-energy maps, i.e. the explored regions. The position of the clusters of the free-energy maps are quite similar among REMD and ST, but each simulation stays a different time in each cluster. To have a more quantitative analysis of the *inter-protocol* agreement, we traced the evolution of RMSD profile-profile JSD distances comparing both protocols, as we compared the runs inside a single protocol (*intra-protocol* convergence). Therefore we merged all available runs into one large trajectory per protocol and took the same number of frames per protocol by sub-sampling the trajectory with a higher number of frames. With the data from each protocol we generated an average RMSD profile which each protocol should reproduce. The JSD curves in figure 5 show that four peptides (7A, 7C, 8A, 8B) showed a good agreement among the REMD and ST protocols, whereas the others, mainly larger peptides, do not. For the cases with a lower agreement (7B, 8C, 9A, 10A, 10B) the curves are almost horizontal at the second half of the trajectories, indicating that longer simulation times would not help here to obtain a better agreement among both protocols.

**Figure 5.**
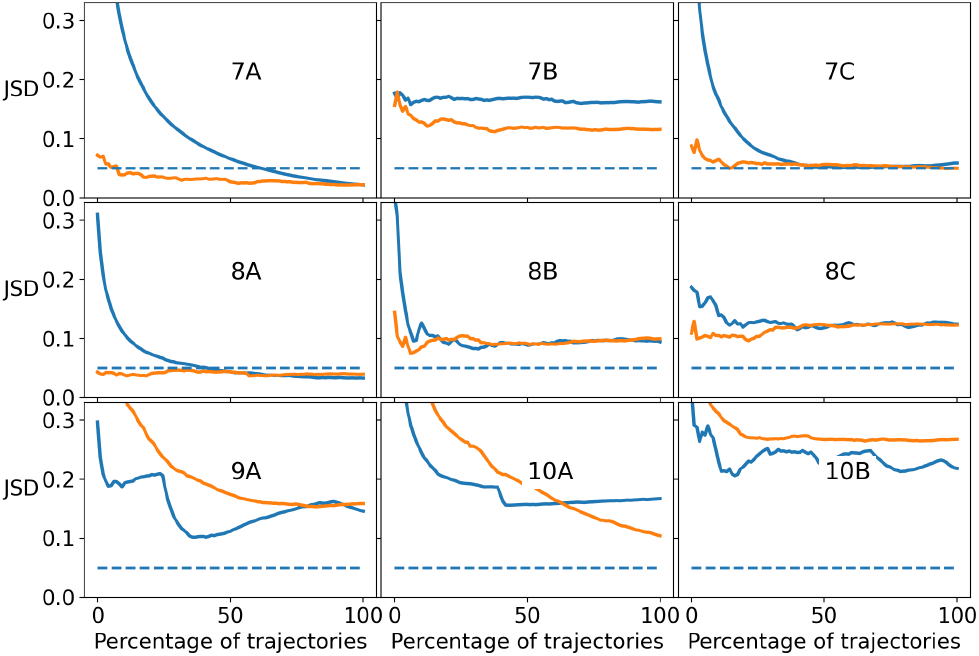
Speed of REMD with ST agreement as measured by the JSD of the RMSD profile of a simulation protocol (ST in blue and REMD in orange) to the averaged RMSD profile (combining ST and REMD profiles). The results obtained with the Amber96 force field are shown here.

To summarize the number of cases where each protocol converges independently and taken together, we calculated the maximum JSD value of the endpoints of each JSD curve, as a metric to assess intra-protocol convergence or interprotocol agreement at the end of the simulated trajectories. The barplot shown in figure 6 confirms that convergence is obtained more often by simply repeating a simulation with the same protocol (REMD or ST here) than by doing a simulation with different MD simulation methods, as REMD and ST here. Nevertheless, both methods agreed quite well, especially for the six smaller peptides (7A-8C). The three larger peptides (9A, 10A, 10B) challenged intra-protocol convergence and inter-protocol agreement.

**Figure 6.**
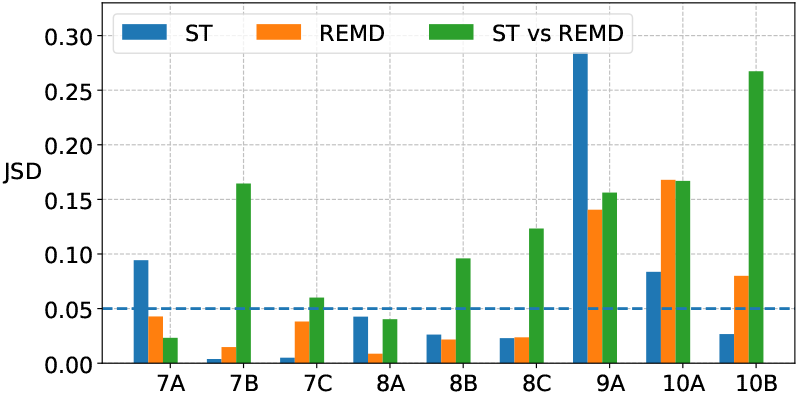
Final convergence or agreement as measured at the end of the trajectories by the maximum JSD of the RMSD profile of a run or simulation protocol to the averaged RMSD profile. The results obtained with the Amber96 force field are shown here. The individual convergence of each simulation protocol (ST in blue and REMD in orange) is compared to the agreement of both taken together (“‘ST vs REMD”‘ in green). The time evolution of the JSD values is shown in figure S2.

For the analyses shown up to here, it has to be reminded, that a single value, as here the maximum JSD value at the endpoint, is quite reductive to describe the similarity of two conformational ensembles. As we will see below, the visual comparison of 2D free energy maps is more informative. As an example, while the peptide 7B showed a high maximum JSD of 0.16 in the comparison ST vs REMD in figure 6, the difference is mainly due to high RMSD clusters, present in the REMD simulations, but not in the ST ones for the Amber96 force field (see figure 9). The highest populated cluster was identical among both simulation protocols. On the other hand, the peptide 9A had the same maximum JSD value as peptide 7B, but the comparison of the free energy maps shows that the REMD and ST maps of peptide 7B for the Amber96 force field look more similar than the ones of the peptide 9A. This visual difference in 2D is not well represented by a single JSD value.

Doing the same analysis as in figure 6 to quantify the inter-protocol and inter-force field agreement for all tested simulation conditions (MD protocol and force field), we obtained the nine heatmaps in figure 7, one for each peptide. The average and standard deviation over all nine peptides are shown in figure 8. The individual heatmaps of each peptide should help in the comparison of the free energy maps shown in figure 9. The order of the columns is the same in figures 7 to 9. The first four columns correspond to the REMD protocols and the last four columns correspond to the ST protocols (as indicated by “‘ST-”‘ before the force field name, absence of it means REMD protocol). For each protocol, one simulation is done with an explicit solvent (columns 4 and 8, as indicated by “‘-exp”‘), whereas the others are done with an implicit solvent. Figure 7 show the singular behaviour of the Charmm36m force field in the ST protocol with the high JSD values in the corresponding “‘ST-Charmm36m”‘ row or column. For the peptide 8A the two REMD simulations in implicit solvent with the Amber14 and the RSFF2C force fields show quite high JSD values to the other simulations, as will be discussed in the next section analysing the free energy maps of figure 9. For the peptide 7C the RMSD profile of the explicit solvent REMD simulation with the RSFF2C force field is only similar to the other explicit solvent simulation in our test set: ST-Amber14-exp, even though the force field and the simulation protocol are different.

**Figure 7.**
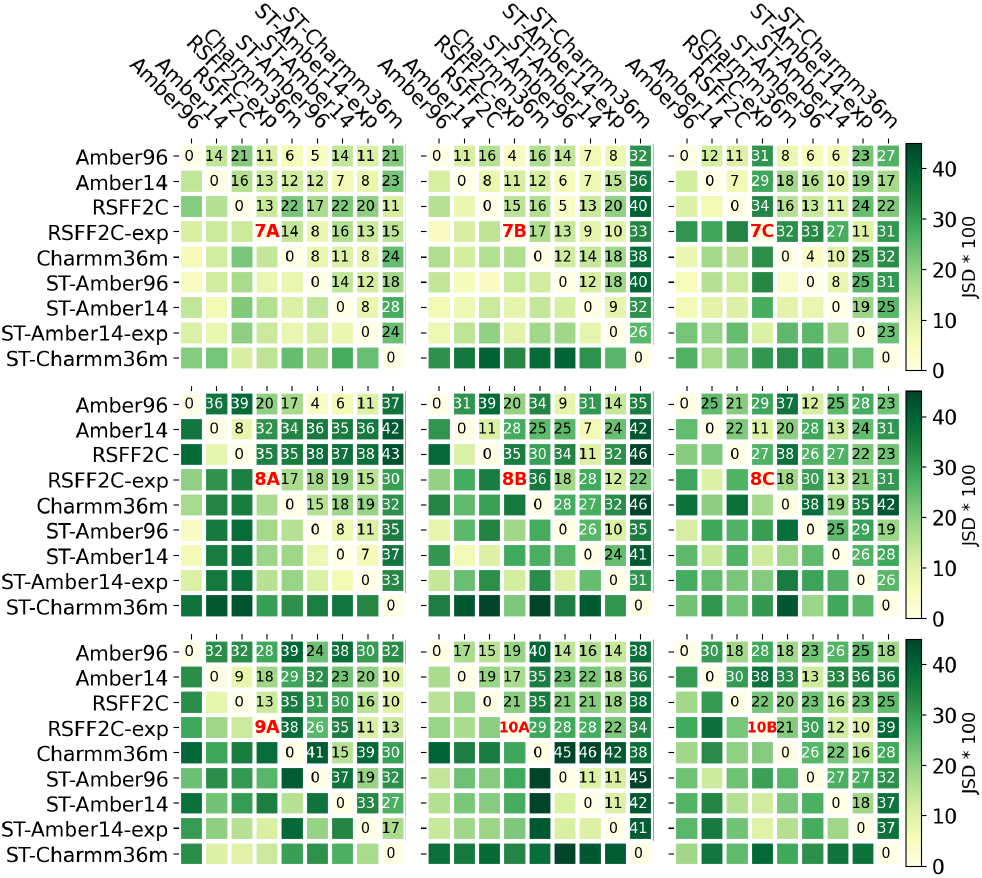
Inter-protocol and force field agreement as measured by the mean JSD of the RMSD profiles to the averaged RMSD profile. The mean JSD is calculated here at the end of the trajectories, as a measure of the agreement obtained at the end of the REMD or ST simulations. Agreement between force fields and simulation protocols (ST where indicated, REMD otherwise). “‘exp”‘ stands for explicit solvent, all other simulations were done in implicit solvent.

**Figure 8.**
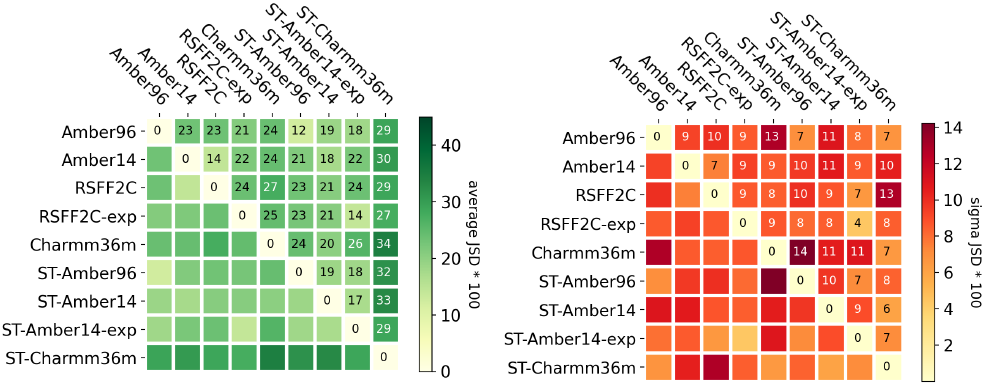
By averaging through the heatmap values of the nine peptides shown in figure 7, the “‘average JSD”‘ values are reported here on the left with their respective standard deviations on the right.

**Figure 9.**
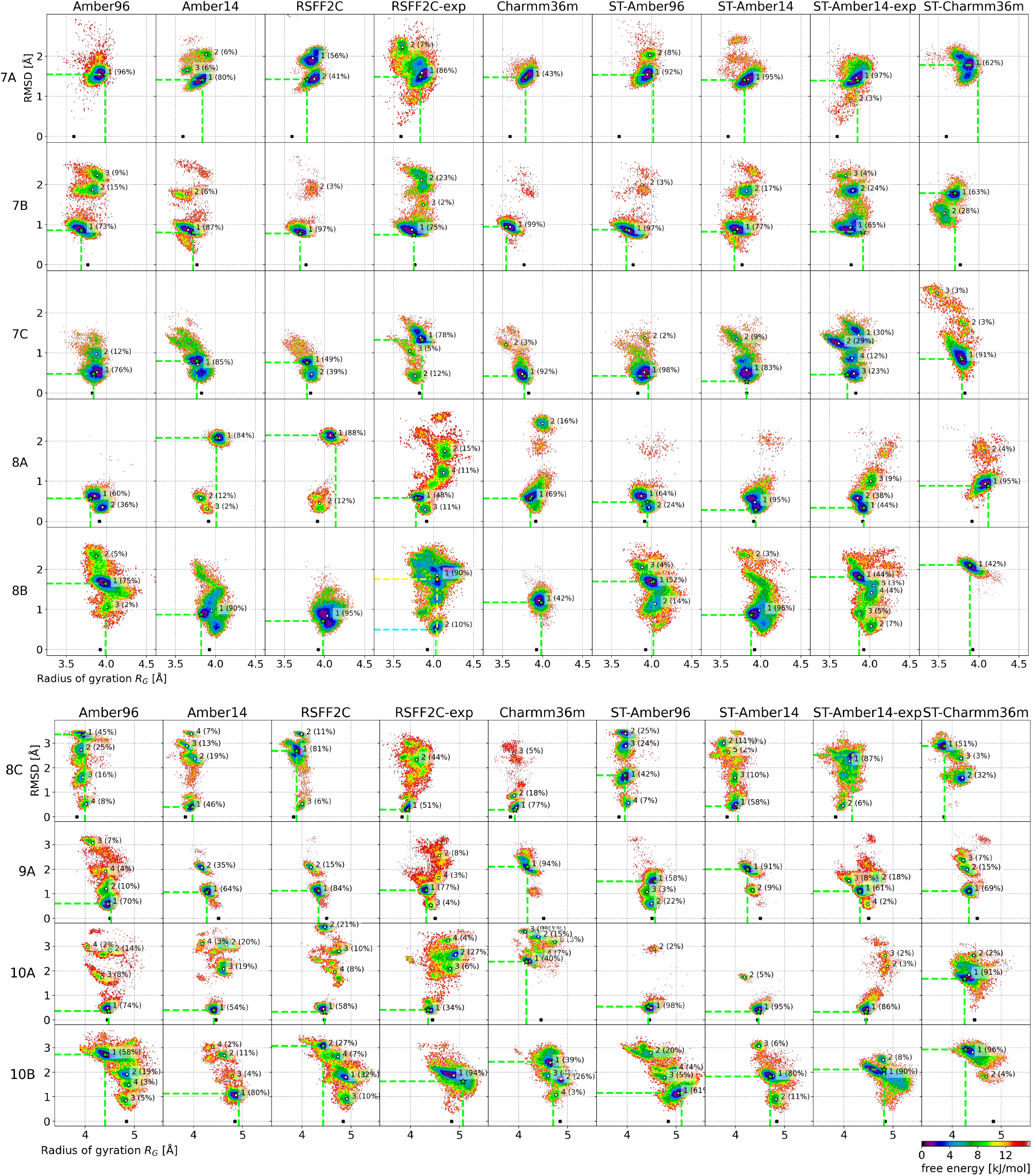
Free energy maps projected onto the radius of gyration (horizontal axis) and the RMSD to the reference structure. The units of the axes are in Å and the free energy unit is in kJ/mol. The first five columns were obtained with the REMD protocol and the last four columns with the ST protocol. The force field employed is indicated on top of each column. RSFF2C-exp and ST-Amber14-exp are explicit solvent simulations. The other columns are implicit solvent simulations. The clustering results are indicated by the cluster ids and their fraction of the total number of frames, from the most populated cluster (id = 1) to the less populated cluster (clusters with a size below 2% are not shown). The centroid of each cluster is indicated by white dots. The centroid position of the most populated and lowest free energy cluster of the PCA projection (see Figure S4) is indicated by a green star and green dashed lines. In the rare case that the position of the lowest free energy value in the PCA maps is different from the position of the most populated cluster, we plotted a cyan star for the former and a yellow star for the latter. The radius of gyration of the reference experimental structure is indicated by a black cross on the 0.0 RMSD line.

Overall figure 8 in comparison to figure 4 confirms that agreement across force fields and/or simulation protocols (i.e. *inter-protocol*) is lower than the convergence of repeated simulations under the same conditions (i.e. *intra-protocol*). The lowest JSD values, i.e. best agreement, are observed for the Amber96 force field, comparing REMD with ST simulation protocol (JSD = 0.12±0.07 (mean±SD)). The fact that the RSFF2C force field is derived from the Amber14 force field is confirmed by the second best JSD value of 0.14±0.07 comparing the REMD simulations with Amber14 and RSFF2C in an implicit solvent. The two explicit solvent simulations, REMD RSFF2C-exp and ST-Amber14-exp, hold the third best JSD value of 0.14±0.04, hinting to the visual similarity of the free energy maps obtained in the two corresponding columns in figure 9.

The switch in MD protocol (REMD↔ST) had overall a lower impact (average JSD values: 0.12±0.07 for Amber96 and 0.18±0.11 for Amber14) than the switch of the force field family from Amber96 to Amber14 (or RSFF2C): average JSD of 0.23±0.09 for REMD and 0.19±0.10 for ST. The switching from implicit to explicit solvent had a greater impact on the RSFF2C force field with REMD than on the Amber14 force field with ST, as quantified by the JSD values of 0.24±0.08 (RSFF2C ↔ RSFF2C-exp) and 0.17±0.09 (ST-Amber14 ↔ ST-Amber14-exp), respectively.

### Free energy maps obtained with REMD and ST on several force fields

Figure 9 shows the free energy maps of seven different simulation conditions over all nine peptides of our dataset. Overall, the three force fields Amber96, Amber14 and the Amber14 variant, RSFF2C, produce similar free energy maps whereas the Charmm36m force field gives quite different results .

The observation of multiple free energy minima in almost each map shows that the cyclic peptides of our dataset have some conformational flexibility, despite the fact that all of them, except peptide 8C, were designed to have a high conformational stability. Only the smallest cyclic peptides of our dataset with seven amino acids (7A-C) show in most cases only one wide free energy minimum. It is difficult to prove that the alternative free energy minima with high RMSD distances to the experimental reference structure exist *in vitro*, as they have not been observed in the NMR studies that produced the reference structures. But some of these alternative free energy minima were observed across several simulation conditions (MD protocols and force fields), which should give a strong evidence for their existence. For example the peptide 8B has a low free energy region centred at a RMSD of 1.7Å and *R*_*G*_ = 4.0Å that is observed in all but two (Amber14 and RSFF2C) simulation conditions. The more flexible peptide 8C has a low free energy “‘band”‘ spanning a large interval of RMSD values from 0.2Å to 3.5Å almost without interruption, i.e. white areas in the maps. Several free energy minima are observed in this “‘band”‘, across all simulation conditions. The larger peptide 10B shows a similar free energy landscape, with the difference that the high RMSD regions are shifted to lower *R*_*G*_ values than the one of the experimental reference structure. Only the two explicit solvent simulations (RSFF2C-exp and ST-Amber14-exp) do not sample significantly the highest RMSD region above 2.8Å and the lowest RMSD region below 1Å, they are both centred in a more restricted RMSD interval of [1, 2.8]Å, compared to the implicit solvent simulations that span a large RMSD interval of [0.5, 3.5]Å. This may be explained by the slowdown of conformational changes of the peptide due to the explicit water molecules, as found already in. ^88^ But for the other ten residues peptide 10A the opposite is observed, when comparing ST-Amber14 with ST-Amber14-exp. The latter has spans to higher RMSD regions.

Looking at the impact of the MD protocol (REMD vs ST), one observes that overall, the maps are quite similar for a given force field, showing that the simulations are well converged and that the sampling method does not influence much the obtained free energy maps. Some small differences can be observed nevertheless: For some peptides, regions with high RMSD conformations compared to the experimental reference structure are more populated by the REMD simulations compared to the ST simulations. This can be seen the best on peptides 9A (Amber 96 only) and 10A (Amber 96 and Amber 14). The lowest free energy cluster is very similar for a given force field, except for some cases where this cluster is lower in RMSD for one method compared to the other. REMD yields a lower RMSD for the peptides 9A and 10B (Amber 14 both) and ST yields a lower RMSD for the peptides 8A (Amber 14), 8C and 10B (Amber 96), see figure 10b.

**Figure 10.**
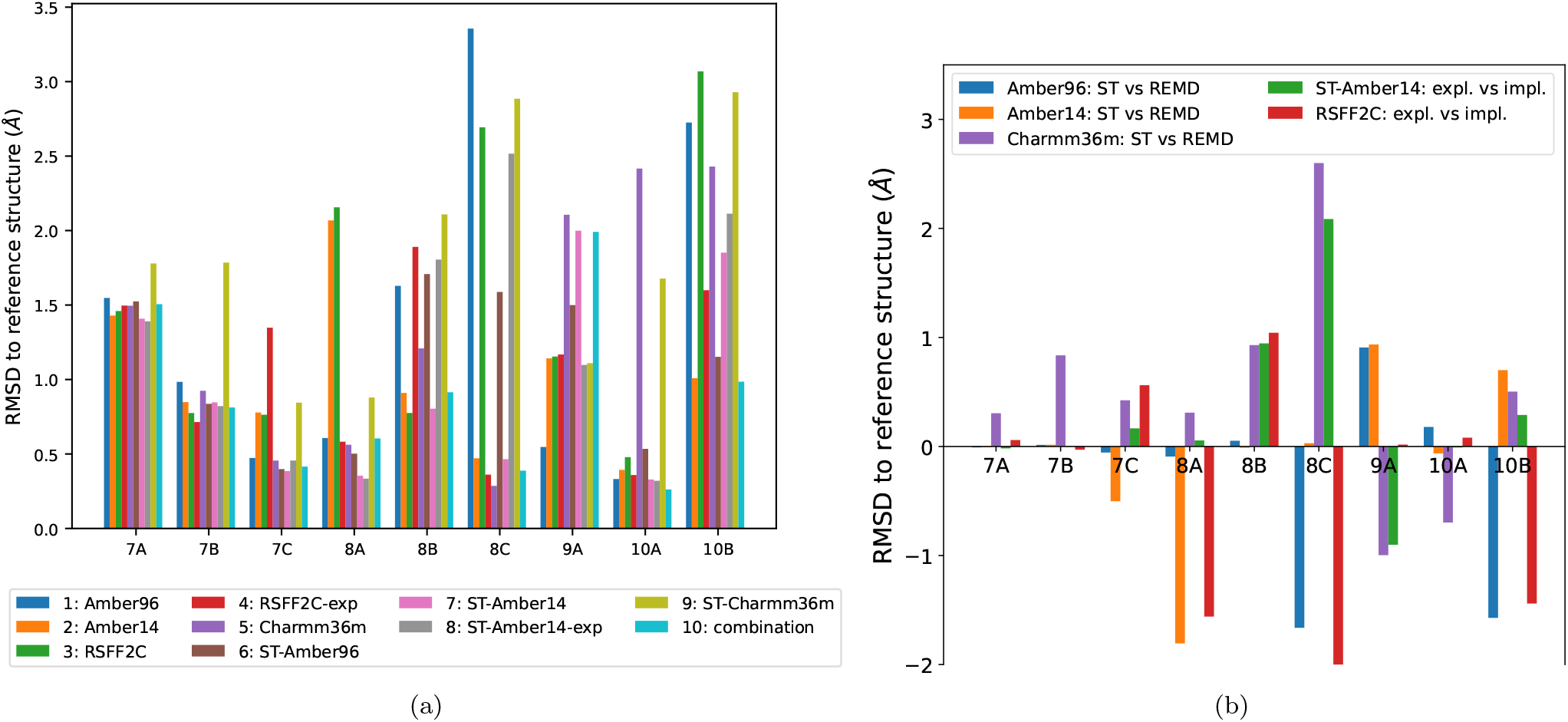
(a) RMSD to the reference structure for the best ranked cluster for each peptide. Results shown for each peptide. The last bar is the combination of Amber96, Amber14, ST-Amber96 and ST-Amber14. (b) Comparison of the two simulation methods, REMD and ST, for three force fields Amber96 (blue), Amber14 (orange) and Charmm36m (purple) in implicit solvent. The RMSD values of the lowest free energy cluster obtained in either method are compared by subtracting the RMSD value obtained by REMD from the one obtained by ST. Positive values indicate cases where REMD give better results and negative values indicate cases where ST give better results. The green bars compare the explicit solvent ST-Amber14 simulation to its implicit solvent equivalent. The red bars compare the explicit solvent REMD-RSFF2C simulation to its implicit solvent equivalent. For the green and red bars positive values indicate cases where the implicit solvent give better results and negative values indicate cases where the explicit solvent give better results.

### Prediction of the experimental reference structure

Applying the protocol described in the materials & methods section to predict the experimental reference structure from our MD simulations we obtained the best clusters indicated by the green stars and green dashed lines in figure 9. The green stars correspond to the most populated and lowest free energy cluster in the PCA maps (figure S4) which are constructed without the knowledge of the experimental reference structure. As can be seen in figure 9 the positions of the green stars in the RMSD/radius of gyration projection superpose very well with the global free energy minima of these maps. As described in the materials & methods section, in the case that the position of the lowest free energy value in the PCA maps is different from the position of the most populated cluster, we plotted a cyan star for the former and a yellow star for the latter. This rare case happened only in one map, namely for the peptide 8B with the RSFF2C-exp force-field: the cyan star gave here a much lower RMSD compared to the yellow star (0.5Å vs 1.8Å). In the further analysis we retained the position of the yellow star for peptide 8B, as this single case cannot validate one prediction method over the other.

In figure 10a we reported the RMSD values of the green (or yellow) stars for each MD simulation. To see the full context of these RMSD values, one should keep a look also on the full free energy maps, shown in figure 9. We remind here that the experimental reference structure is here the best representative conformer of the NMR ensemble (first model here). In addition, we calculated also the minimum RMSD of the best ranked cluster to the 20 NMR models of each peptide, see figure S5. As this only reduces globally the RMSD values without changing the profiles of 10a, we kept here the RMSD values to the best representative conformer of the NMR ensemble.

For the smallest cyclic peptides of the set, with seven amino acids (7A-7C), the different simulations converged to similar RMSD values (figure 10a). Comparing the RMSD values among these three peptides 7A to 7C, one can observe that the peptide 7A has a threefold higher RMSD than the peptide 7C (1.5Å vs 0.5Å). This shows that the notion of “‘low”‘ RMSD is specific to each peptide.

For the other peptides, only the peptide 10A shows a very good agreement among the force fields and simulation protocols: All simulations, except the two using Charmm36m, yield the same global free energy minimum at only 0.4 - 0.5Å (figure 10a), despite the relatively large size of the peptide 10A of 10 amino acids, that allows a large range of conformations with up to 4Å of RMSD to the reference structure (figure 9).

The performance on the peptide 10A is even more impressive, if it is compared to the second peptide with 10 amino acids in the set: the peptide 10B. Here the lowest RSMD is only 1.0Å obtained with two simulations: ST-Amber96 and REMD Amber14 (figure 10a). Nevertheless, all simulations sampled with a significant frequency the sub 1Å region (see figure 9). The differences among the simulations in the population sizes of the clusters may come from insufficient convergence, as REMD and ST show here quite different results, even if the same force field is used (figure 10b). It may come also simply from a relatively flat free energy landscape for the peptide 10B, that does not give a clear preference for the experimental reference conformation.

The results obtained on the peptide 8A are quite similar to the peptide 7C, except for the REMD Amber14 and RSFF2C simulations in implicit solvent. The other seven simulations of peptide 8A give a very low RMSD of 0.4−0.6Å (figure 10a) and do not sample much the higher RMSD regions (figure 9). The REMD Amber14 and RSSF2C simulations sample the same low RMSD regions as the other simulations, but at a lower frequency than the unique cluster at 2.2Å (figure 9), that is not sampled at all by the other simulations. We investigated in deep this high RMSD cluster of the peptide 8A, see supplementary section 8.

For the peptide 8B the Amber14 force field and its derivative RSFF2C give the best results in terms of low RMSD with a good value of 0.9Å. The use of an explicit solvent degraded this good result here for both force fields (figure 10b). Here the two simulation protocols, REMD and ST, give exactly the same results (figure 10b), also the free energy maps look very similar (figure 9).

The peptide 8C is particular as it contains only L-amino acids and is therefore probably more flexible than the other peptides that contain a mixture of L and D-amino acids. In fact, the free energy maps of 8C show a large landscape from 0.5Å to 3.5Å of RMSD (figure 9). For such a flexible peptide it is difficult to rank the lowest RMSD cluster in the first position. The Amber14 force field pick up the right cluster, with a very low RMSD of 0.4Å. The use of an explicit solvent degrades this performance, as the ST-Amber14 explicit solvent simulations gives a high RMSD cluster of 2.5Å as the most populated cluster. Interestingly, for the RSFF2C force field it is the opposite case (figure 10b): the explicit solvent improved the ranking with the lowest observed RMSD for 8C of 0.3Å, instead of a high RMSD of 2.7Å for the implicit solvent simulation. The use of the Amber96 force field gives here even worse results, with 3.4Å using the REMD protocol. The ST protocol yields also a quite low free energy basin at 3.4Å, but the basin at 1.5Å has here a slightly lower free energy, explaining the great difference of the two protocols using the Amber96 force field for the peptide 8C (figure 10a). Both protocols result here in a quite similar free energy map (figure 9, peptide 8C), showing that the differences observed in figure 10a should not be over-interpreted, a look on the full free energy maps is still necessary.

The last peptide to be analyzed is the peptide 9A: Here the REMD Amber96 simulation gives the best results of 0.6Å. The Amber14 force field give a doubled RMSD value (1.2Å) for the REMD protocol and for the ST protocol, but only in explicit solvent. The ST-Amber14 implicit solvent simulation ranks the higher RMSD cluster of 2Å at the first position. The RSFF2C force field give almost identical free energy maps as with Amber14 force field using the REMD protocol. The explicit solvent RSFF2C simulation populates the 0.6Å cluster a bit more than its implicit solvent counterpart (figure 9).

### Explicit vs implicit solvent

The Amber14 force field has been optimized for the use of an explicit solvent, therefore we expected better results using this force force field with explicit solvent than with implicit solvent. Especially the comparison of the ST-Amber14 implicit solvent simulations with its explicit solvent counterpart (ST-Amber14-exp column in figure 9) should show the improvements by the use of an explicit solvent vs an implicit solvent. But surprisingly this is observed only for the peptides 7A (lowest free energy cluster does not improve, but lower RMSD areas are explored, see figure 9) and 9A (negative green bar in figure 10b). If the comparison for the peptide 9A is done against the other implicit solvent simulations, the lowest RMSD cluster is as well or better ranked with the three REMD simulations (Amber96, Amber14 and RSFF2C), showing that an explicit solvent is not a requirement to obtain a good ranking. In contrary, we observed more cases where the use of an explicit solvent degrades the ranking for ST-Amber14, namely 8B, 8C and 10B (positive green bars in figure 10b).

### RSFF2C force field

Finally, we turn to looking at the free energy maps obtained with RSFF2C which is a derivative from the Amber14 force field. The MD simulations are done in implicit and explicit solvent using the REMD protocol (RSFF2C and RSFF2C-exp, respectively). For most peptides the results obtained with RSFF2C are quite similar to the results obtained with Amber14 in implicit solvent for both sets, REMD and ST (see figure 9). The peptide 8A is a special case, as only the two REMD simulations with Amber14 and RSFF2C found a large cluster at a high RSMD of 2.2Å. This special case underlines the similarity of the two forces fields Amber 14 and RSFF2C, at least in implicit solvent conditions. The RSFF2C simulations in explicit solvent (RSFF2C-exp) gave better results for three peptides (8A, 8C and 10B, see negative red bars in figure 10b) and gave only for two peptides, 7C and 8B (positive red bars in figure 10b), worse results, compared to the implicit solvent simulation (RSFF2C). This is the opposite result for what we found when comparing implicit vs explicit solvent for the Amber14 force field (see above), despite the fact that the RSFF2C force field is derived from the Amber14 force field. It seems therefore more important to have an explicit solvent with the RSFF2C force field, than with the Amber14 force field.

### Combining different force fields and simulation protocols

As described in the methods section, we combined different simulations to achieve more robust results, especially in the prediction of the experimental reference structure. The rationale behind this approach is the observation that while the low RMSD clusters are often the same among the different simulations, high RMSD clusters are more spread. Combining the simulations should increase the populations of the low RMSD clusters, while decreasing the ones of high RMSD clusters, and therefore increasing the chance to rank the low RMSD clusters at the first position. As we have already seen in figure 10a none of the simulations gives a good prediction for all nine peptides of our dataset. This can also be seen in figure 11a, where the RMSD values of the nine peptides shown in figure 10a are assembled into a box plot per simulation. The upper limit extends often above 2.0Å. The single simulations are sorted here by the median RMSD of the nine RMSD values of each peptide.

**Figure 11.**
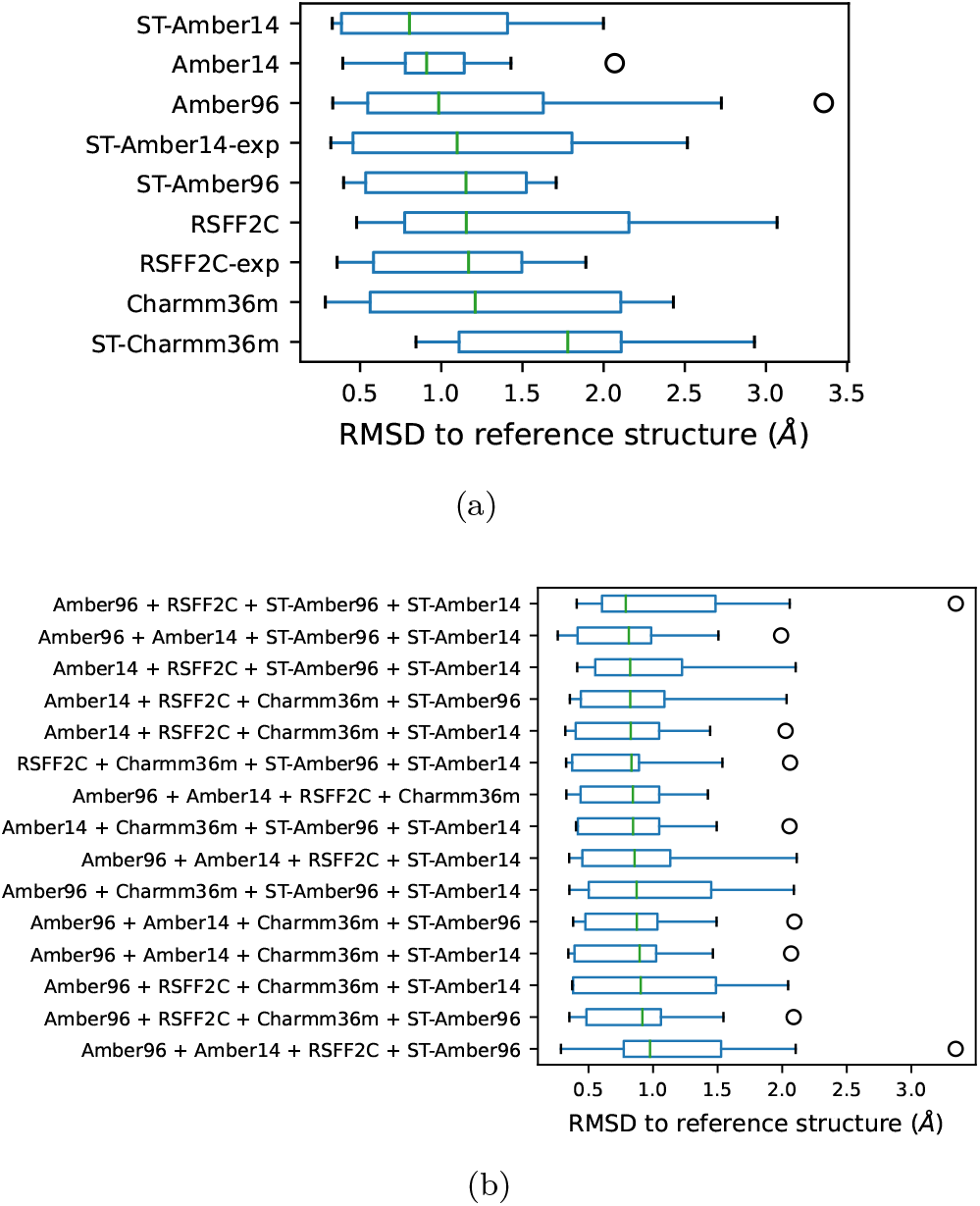
(a) For each simulation the RMSD values of the nine peptides shown in figure 10a are assembled into a box plot. The simulations are sorted here with increasing median RMSD values, shown with green lines. (b) All 15 combinations of four implicit solvent simulations without ST-Charmm36m.

The median RMSD values of figure 11a are reported on the first column in figure 12a. If only a second simulation is added to one of the nine simulations, the median RMSD values are significantly improved, as can be seen by comparing the violin plots of the first two columns in figure 12a. Figure S18 gives the details of the 36 possible combinations of twins among the nine simulations (binomial coefficient: 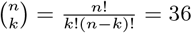 with *n* = 9 and *k* = 2 here).

**Figure 12.**
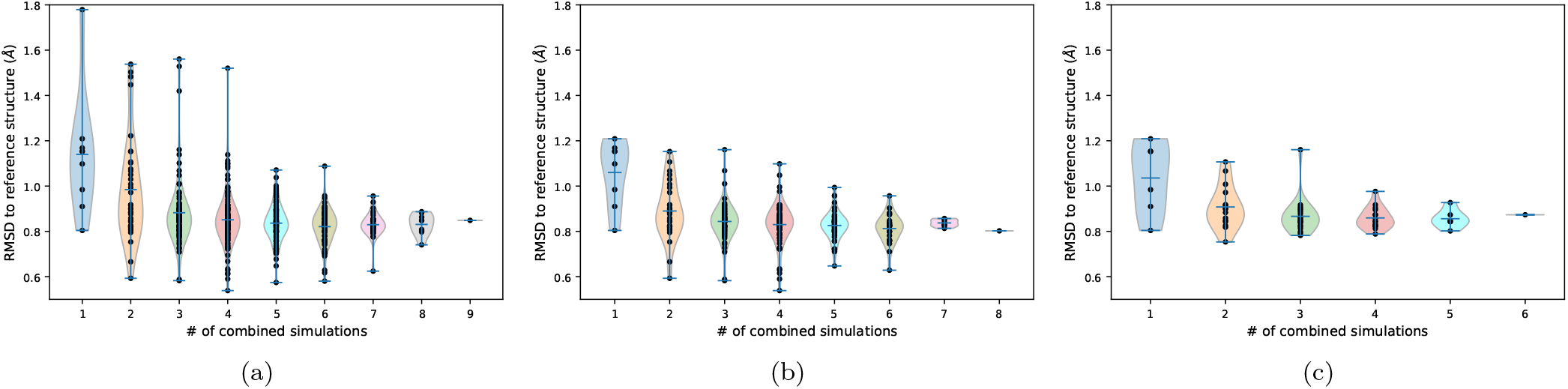
Combining simulations to improve the predictions of the reference structure. See supplementary figures for more details. (a) Predictions for all possible combinations of simulations. One point corresponds to one simulation (first column), two combined simulations (second column) or any higher number of combined simulations (columns three to nine). The y-value of each point is the median RMSD to the reference structure calculated over all nine peptides. The violin-plots show the distribution of the points along the y-axis. (b) Same plot as in (a), but without the ST-Charmm36m simulation. (c) Same plot as in (a), but without the ST-Charmm36m simulation and with implicit solvent simulations only.

Looking at the median RMSD values in figure S18, represented by green bars, the top 3 twin combinations are: Amber96 + RSFF2C-exp, RSFF2C-exp + ST-Amber14 and RSFF2C + ST-Amber14. As many twin combinations are clustered around 0.9Å for the median RMSD (see also figure 12a), other criteria might be used in addition to rank or select twin combinations. For example the upper and lower limits of the box plots in figure S18. Here the Amber14 + ST-Amber14 twin shows a good compromise with a low upper limit and a not too high lower limit in the RMSD scale. But due to the high number of 36 twin combinations and the similarity of the values, this selection or the one of the top 3 should not be used as a general rule. What is more important is the result that the combination of two simulations is for the majority better than a single simulation taken alone. Combining more than two simulations continues to improve the RMSD distributions in figure 12a, especially by reducing the upper limit of these distributions. Combining all nine simulations yield a median RMSD as low as the best single simulation (here ST-Amber14). As in another set of peptides the ST-Amber14 might not be the best simulation, the combination of diverse simulations yields robustness in the choice of the right simulations.

Instead of trying to identify the single best simulation among our set of simulations, we can exclude the worst simulation, here ST-Charmm36m which has an important gap in median RMSD to the other eight simulations (see figure 11a). Unsurprisingly this improves the RMSD distributions for all simulation combinations (see figure 12b).

Producing eight REMD/ST simulation trajectories per peptide is quite costly, especially for the two explicit solvent simulations. Fortunately, discarding both in addition to ST-Charmm36m does not degrade significantly the RMSD distributions (see figure 12c). There are essentially less lowest and highest RMSD values, probably due to the reduced number of combinations. For example for four combined simulations, there are 70 combinations among eight possible simulations (figure 12b) but only 15 combinations among six possible simulations (12c). Producing four implicit solvent REMD and ST simulations may be a good compromise in terms of robustness vs calculation cost, as the median RMSD values are all below 1.0Å.

All 15 combinations of implicit solvent simulations are listed in figure 11b in the order of increasing median RMSD values. The lower and upper bounds of the second best are lower than the combination ranked in first position. Despite its slightly higher median RMSD value, it is one of the best choices among the 15 combinations. The second best combination is: Amber96, Amber14, ST-Amber96 and ST-Amber14. The combination at the 7th position is composed of only REMD simulations: Amber96, Amber14, RSFF2C and Charmm36m. It yields an even lower upper bound and may be a viable choice to reduce the time to setup two different simulation protocols, here REMD and ST. Even if RSFF2C and Charmm36m are lower ranked as single simulations (see figure 11a) than ST-Amber96 and ST-Amber14, they nevertheless enrich the diversity in free energy maps, necessary to obtain robust predictions of the reference structure.

Overall, this shows that a limited number of implicit solvent simulations with the two Amber force fields, Amber96 and Amber14, and with the two simulation methods, REMD and ST yield already robust results, predicting the experimental reference structure accurately for 8 out of 9 cyclic peptides. Only for the peptide 9A the best cluster of the Amber96 + Amber14 + ST-Amber96 + ST-Amber14 combination is at a higher RMSD of 2.0Å, while the other eight peptides are below 1.0Å or at 1.5Å for the peptide 7A. See the last cyan bars of “‘10: combination”‘ in figure 10a. Just looking at the RMSD values of the selected best clusters in figure 10a does not explain the high RMSD value for peptide 9A. It has to be reminded here that the RMSD values of the four combined simulations are not obtained by simple averaging over the RMSD values of the four individual simulations, but by combining the MD trajectories and from the PCA analysis of the combined trajectory. Looking at the free energy maps of the individual and combined simulations gives a clearer picture, see figure S6. The two Amber14 simulations (REMD + ST) have an important population on the cluster of 2.0Å to the reference structure, especially for ST with 91%. The two Amber96 simulations (REMD + ST) show both a cluster at low RMSD, especially for REMD with 70%, but this is still lower than for the high RMSD cluster. At the end the high RMSD cluster has 31% and the low RMSD cluster 21% in the combination of the four simulations. But at least the difference in population between both clusters is much lower than for both Amber14 simulations, where the the same low RMSD cluster is almost missing completely.

Finally, we would like to mention that as the peptides are not simulated in interaction with a binding partner, there is no need here to use explicit solvent simulations to generate accurate free energy landscapes. Three or four implicit solvent simulations will take less computational time than a single explicit solvent simulation and more crucially will give robust and accurate results, which is not achieved with a single explicit solvent simulation.

### Validation on four additional peptides

We selected four additional peptides as test set with the PDB-codes: 6UD9, 6UCX, 6UDZ and 6UDW. The peptides listed in table S1 are of the same nature as the nine initial peptides of the study, as they have the same number of amino acids, two of 8 A.A. and two of 10 A.A. and as they have a mixed chirality. All of them were designed peptides from the David Baker group, ^89^ but solved by X-ray instead of NMR.

We applied our recommendation to use the four simulations, Amber96 + Amber14, ST-Amber96 and ST-Amber14, combined. We produced 1000 ns for each of the eight replica of the REMD simulations and 10 *μs* for each ST simulation. The figure S7 shows the 16 free energy maps of each simulation at 300K, as well as the combined free energy maps for each of the four peptides. One can observe that the two REMD simulations had difficulties to explore the low RMSD region for 6UD9 and 6UDW, but not for 6UDZ nor for the Amber14 simulation of 6UCX. ST performed here better, except for 6UCX with the Amber96 force field. The ST-Amber14 simulations performed well, but populated an higher RMSD cluster at the first position for 6UDW. The RMSD values of the best ranked clusters are reported in figure S8. The combination of all four simulations is still beneficial, as the low RMSD region is populated for every peptide, see figure S7. The low RMSD population is highest for three of the four peptides, only for 6UDW a higher RMSD cluster is populated more (42%) than the lowest RMSD cluster (17%).

This validation of the combined use of four simulations shows that even if only half of the simulations have a cluster with a low RMSD, the combined simulation can still recover it, as was seen here on the two peptides 6UD9 and 6UCX. This stems from the fact that the high RMSD clusters are often different in their 3D structures and are therefore not clustered together in the PCA maps (figure S9).

## Conclusion

Our study explored various molecular dynamics “‘ingredients”‘ for the cartography of the conformational landscape of cyclic peptides and the prediction of their reference experimental structures. Three types of ingredients were investigated: the force field, the solvent model and the simulation method. For the force fields, no single best force field could be observed, older force fields like Amber96 and Amber14 performed as well as the recent RSFF2C force field. The use of an explicit solvent instead of an implicit solvent showed a mixed picture: for RSFF2C the prediction of the native structure worked better in explicit solvent, while for Amber14 we observed the opposite. Under the same force field and solvent model both simulation methods, REMD and ST, give very similar free energy maps, especially for the lowest free energy clusters. Some differences could be observed in higher free energy clusters and REMD tended to produce more high free energy clusters than ST. No clear advantage for one method vs. the other in the prediction of the native structure could be observed. The *intra-protocol* convergence of repeated simulations under the same conditions has been observed to be higher than the *inter-protocol* agreement of simulations using different conditions, i.e. force field, solvent model or simulation method. This finding is important to underline, as convergence of simulations is often only evaluated by repeating simulations under the same conditions, which seems to be an insufficient convergence criteria.

Among the nine peptides, the native structure is predicted with a median backbone RMSD ranging from 0.8 to 1.8Å with a single MD simulation The ST-Amber14 (implicit solvent) simulation yielded the best result here. To improve these values we combined systematically all nine simulations, merging two, three or more simulations together. The average median RMSD is gradually reduced from 1.15 to 0.85Å with the number of merged simulations. More importantly, the maximum median RMSD is also greatly reduced from 1.8 to 0.85Å This can be explained by the observation that high RMSD clusters are less conserved among the seven simulations than the lowest RMSD cluster. The merging of the simulations enforces therefore the populations of the lowest RMSD cluster which becomes the most populated cluster more often. The two explicit solvent simulations performed here, are not necessary to obtain good predictions, the six implicit solvent simulations are sufficient for this. As the ST-Charmm36m simulation in implicit solvent had the highest median RMSD of 1.8Å with an important gap of 0.6Å to the next best simulation, we excluded it. Among the 15 combinations of four simulations in implicit solvent, one combination gave a good median RMSD of 0.85Å with a maximum RMSD below 2.0Å: REMD Amber96 + Amber14 and ST Amber96 + Amber14. With a slightly higher median RMSD of 0.9Å, but with a maximum RMSD of only 1.5Å, the combination of four REMD simulations is also an interesting alternative: Amber96 + Amber14 + RSFF2C + Charmm36m.

The combination of different “‘ingredients”‘ of MD simulations, here the force field and the simulation method, seems to yield more robust results, with a better native structure prediction performance. Three or four implicit solvent simulations are already sufficient and will take less computational time than a single explicit solvent simulation. In addition to the combination of complementary MD simulations, our native structure prediction method is based on a PCA projection of the free energy landscape and robustly yield good predictions for the nine peptides. It would be interesting to apply our method also to the structure prediction of proteins and protein-peptide complexes, as well as cyclic peptides with chemical modifications, like N-methylation.^35^

## Data and Software Availability

ST simulation protocol:

https://github.com/samuelmurail/SST2/tree/main/bin

Trajectory data and python scripts for analysis:

https://doi.org/10.5281/zenodo.15112951

## Acknowledgement

We would like to thank Fan Jiang and Wei Kang from Peking University, Beijing, for sending us their RSFF2C force field parameters files. This project was provided with computer and storage resources by GENCI at CINES thanks to the grant 2021-A0100707641 on the supercomputer Occigen. The RPBS and iPOP-UP platforms also contributed with calculation time.

## Supporting Information Available

Supplementary materials and methods and supplementary figures are available for download.

## Notes

https://doi.org/10.5281/zenodo.15112950

https://github.com/samuelmurail/SST2/tree/main/bin

### Competing Interest Statement

The authors have declared no competing interest.

### Summary of Updates

- Additional REMD and ST data with Charmm36m - Figures 7 to 12 changed, new Figure 11 - Section "Combining different force fields and simulation protocols" changed significantly due to the removal of the RMSD threshold notion. - Approach validated on four additional peptides (see new section "Validation on four additional peptides") - Supporting information (SI): - new sections 5 to 9. - large section 8 on the deep investigations on the special case of peptide 8A

