## Supporting Information for "Robust conformational space exploration of cyclic peptides by combining different MD protocols and force fields"

July 7, 2025

### Contents

**1 Supplementary materials and methods** **3**

1.1 Explicit solvent REMD protocol . . . 3

1.2 Clustering algorithms . . . 3

**2 Convergence of REMD simulations** **5**

**3 ST convergence for the Amber96 force field** **7**

**4 Free energy maps with PCA** **9**

**5 Minimum RMSD among all 20 NMR models** **10**

**6 Combination of four simulations for peptide 9A** **10**

**7 Additional test set of four peptides** **11**

**References** **34**

### 1 Supplementary materials and methods

#### 1.1 Explicit solvent REMD protocol

Temperatures list for the explicit solvent REMD protocol with 32 replicas, as obtained with the web-server https://virtualchemistry.org/remd-temperature-generator/:

300.00, 304.31, 308.67, 313.07, 317.52, 322.01, 326.55, 331.14, 335.78, 340.44, 345.17, 349.96, 354.80, 359.69, 364.63, 369.62, 374.67, 379.77, 384.92, 390.13, 395.39, 400.72, 406.10, 411.53, 417.03, 422.58, 428.19, 433.87, 439.60, 445.39, 451.25, 455.50K.

#### 1.2 Clustering algorithms

For the identification of low free energy clusters we employed two techniques: k-means clustering[1] and HDBscan clustering.[2] The k-means clustering is relatively easy to control manually by adjusting the number of clusters to search. It is a good choice for the clustering of a limited number of 2D projections. For a larger number of 2D maps, a clustering algorithm that does not need a manually adjusted parameter, like the number of clusters of the k-means algorithm, is required. The HDBscan algorithm automatically adjusts the number of clusters. But we had to adjust the "min_cluster_size" and especially the "min_samples" parameters to find the two values that give a good compromise for all maps. For landscapes where the local free energy minima are not well separated by low density regions, they are often merged together by HDBscan. As we searched for a clustering with a number of clusters similar to the number of free energy minima, we had to find a minSamples value that does the right level of "smoothing" of the map, so that HDBscan does not merge too much. Finally we set min_cluster_size to 100 and min_samples to 20 for figure 9 of the article. In addition, as some maps have only one cluster, we activated the "allow_single_cluster" option, otherwise HDBscan will try to find more than one cluster, and possibly give no cluster at all.

For the 2D maps of figure 9 the input to HDBscan is a two columns table with the radius of gyration and RMSD to experimental reference structure for each frame of the trajectory. The same input is fed to the k-means algorithm, employed in figure S1. For the 2D maps of the PCA (figure S4), the input to HDBscan is a multi-column table with the values of the principal components for each frame, in general four to six columns.

For the k-means clustering we employed the "deeptime" python package[3] and for the

HDBscan clustering the "HDBscan" python package.[2]

### 2 Convergence of REMD simulations

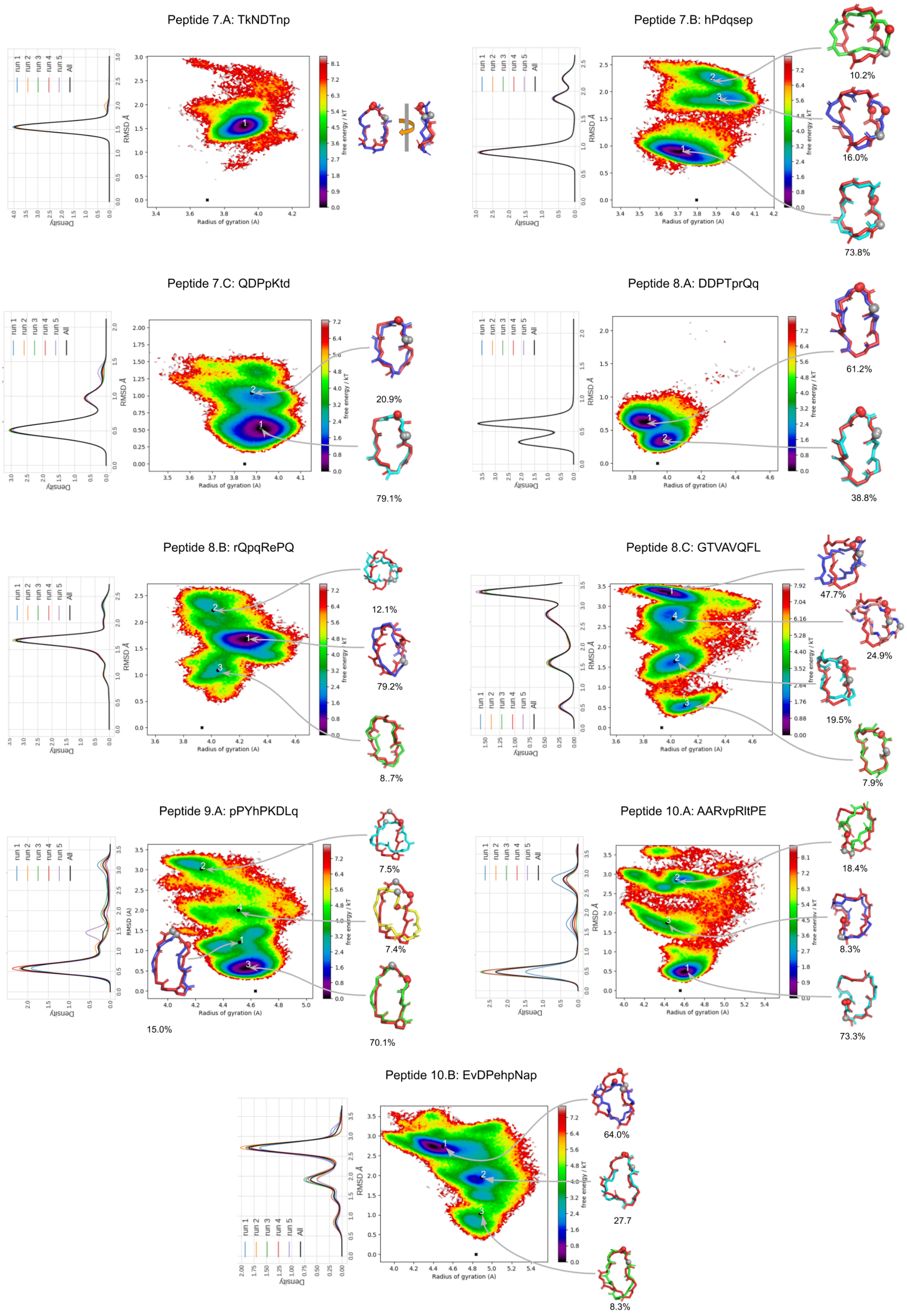

Figure S1: Convergence of simulations assessed by repeating simulations. All results shown in this figure were obtained with the Amber96 force field in implicit solvent using the REMD protocol. For each of the five runs the density profile of the RMSD to the experimental reference structure is shown on the left. The density profile of the five concatenated trajectories is shown in black. The free energy map is also obtained by the assembly of all five runs. The map is clustered with the k-means algorithm and the centroid structures are superposed to the experimental reference structure (in red). The two first residues are indicated with a red and a gray ball. The size of each cluster is indicated below the 3D structures

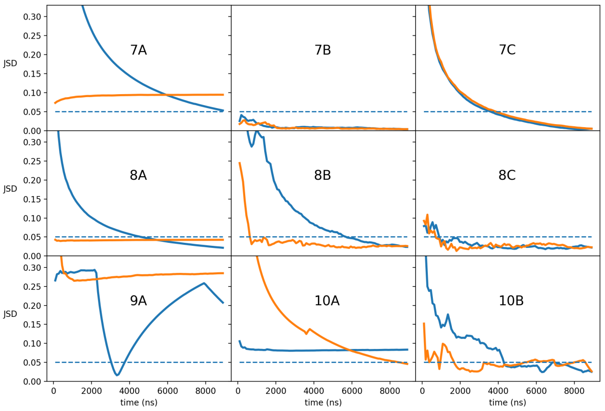

(a)

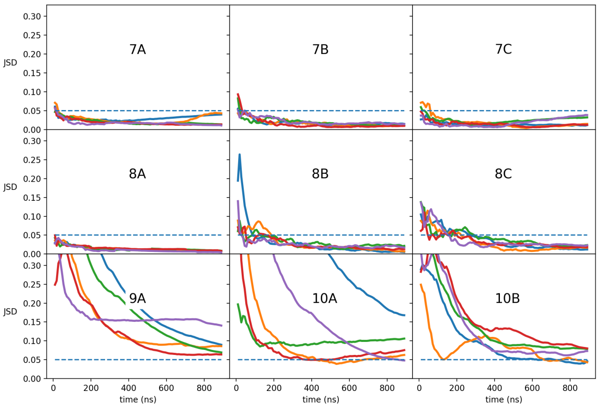

(b)

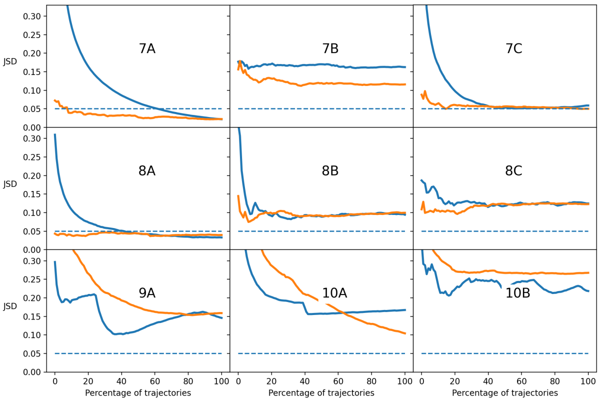

(c)

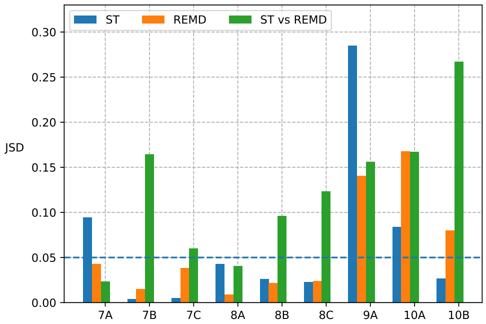

(d)

Figure S2: Speed of convergence as measured by the JSD of the RMSD profile of a run or simulation protocol to the averaged RMSD profile. The results obtained with the Amber96 force field are shown here. Below the threshold of a JSD of 0.05 two RMSD profiles are nearly identical. Convergence is obtained, if all curves fall and stay below this threshold. (a) Two ST runs with the Amber96 force field. Run 1 in blue and run 2 in orange. (b) Five REMD runs with the Amber96 force field. Each color correspond to one run. (c) Convergence of ST (blue) with REMD (orange) with the Amber96 force field. (d) Maximum JSD values at the end of the trajectories, from the three figures a) to c).

### 3 ST convergence for the Amber96 force field

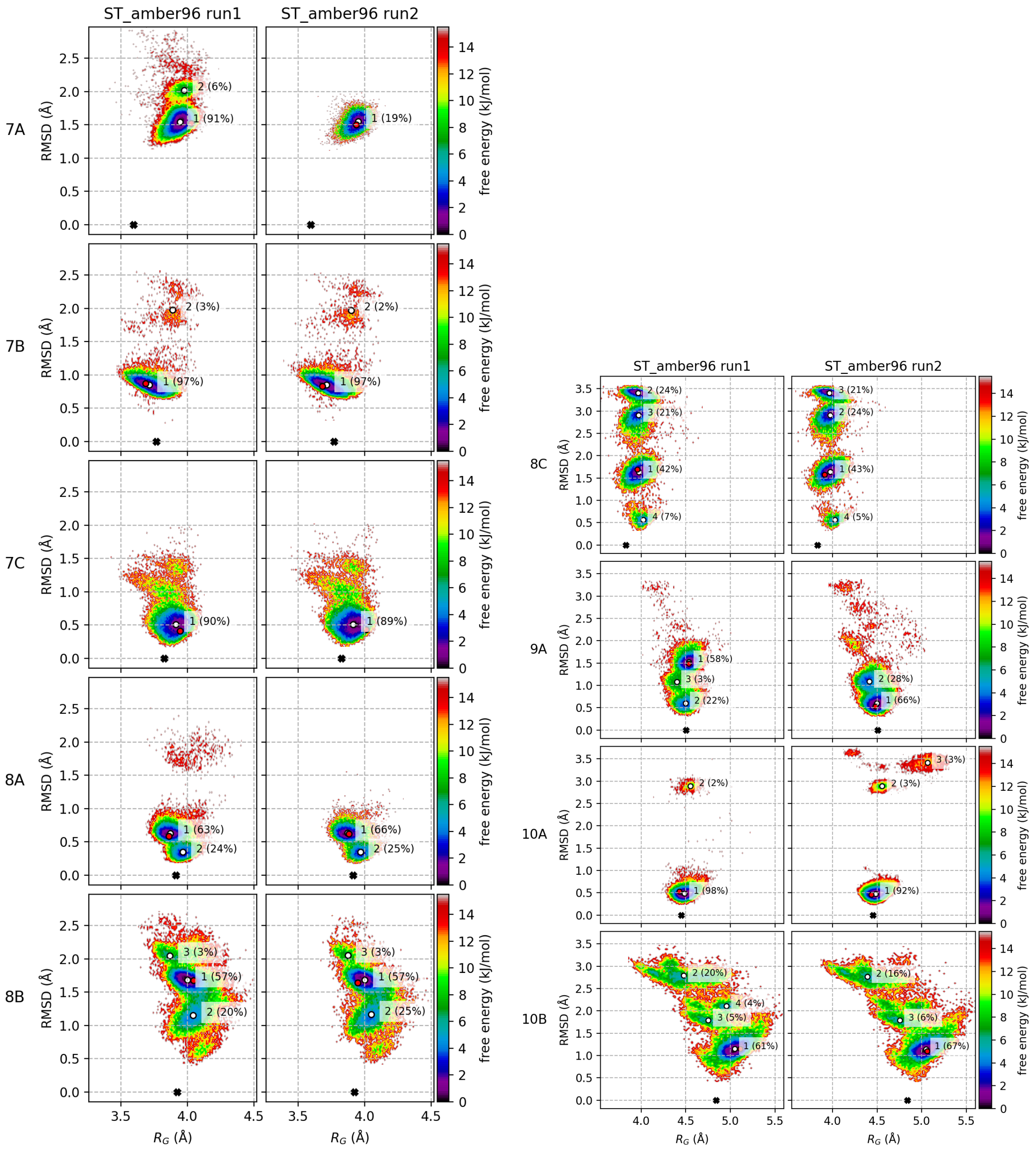

Figure S3: Free energy maps projected onto the radius of gyration (horizontal axis) and the RMSD to the reference structure. The units of the axes are in Å and the free energy unit is in kJ/mol. Both columns are obtained with the ST protocol using the Amber96 force field in implicit solvent, they are just two different runs to test the convergence of ST. The HDBscan clustering results are indicated by the cluster ids and their fraction of the total number of frames, from the most populated cluster (id = 1) to the less populated cluster (clusters with a size below 2% are not shown). The centroid of each cluster is indicated by white dots. The centroid position of the most populated cluster of the PCA projection (not shown) is indicated by a red dot. The radius of gyration of the reference experimental structure is indicated by a black cross on the 0.0 RMSD line.

### 4 Free energy maps with PCA

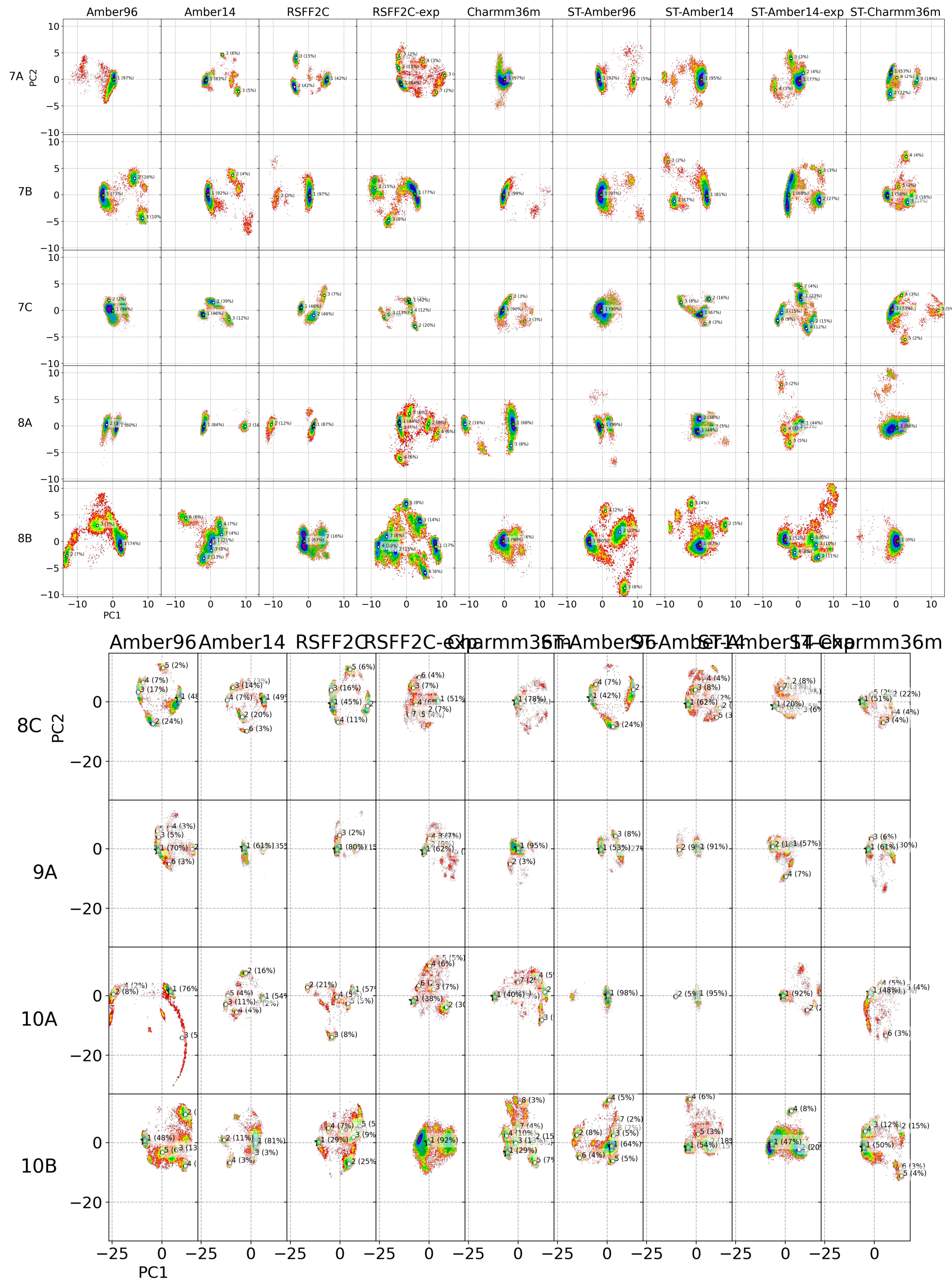

Figure S4: Free energy maps projected onto the first two principal components. The units of the axes are arbitrary and the free energy unit is in kJ/mol. The first five columns were obtained with the REMD protocol and the last four columns with the ST protocol. The force field employed is indicated on top of each column. RSFF2C_explicit and ST_Amber14exp are explicit solvent simulations in REMD and ST, respectively. The other columns are implicit solvent simulations. The HDBscan clustering results are indicated by the cluster ids and their fraction of the total number of frames, from the most populated cluster (id = 1) to the less populated cluster (clusters with a size below 2% are not shown). The centroid of each cluster is indicated by white dots. The centroid position of the most populated and lowest free energy cluster is indicated by a green star. In the rare case that the position of the lowest free energy value is different from the position of the most populated cluster, we plotted a cyan star for the former and a yellow star for the latter.

### 5 Minimum RMSD among all 20 NMR models

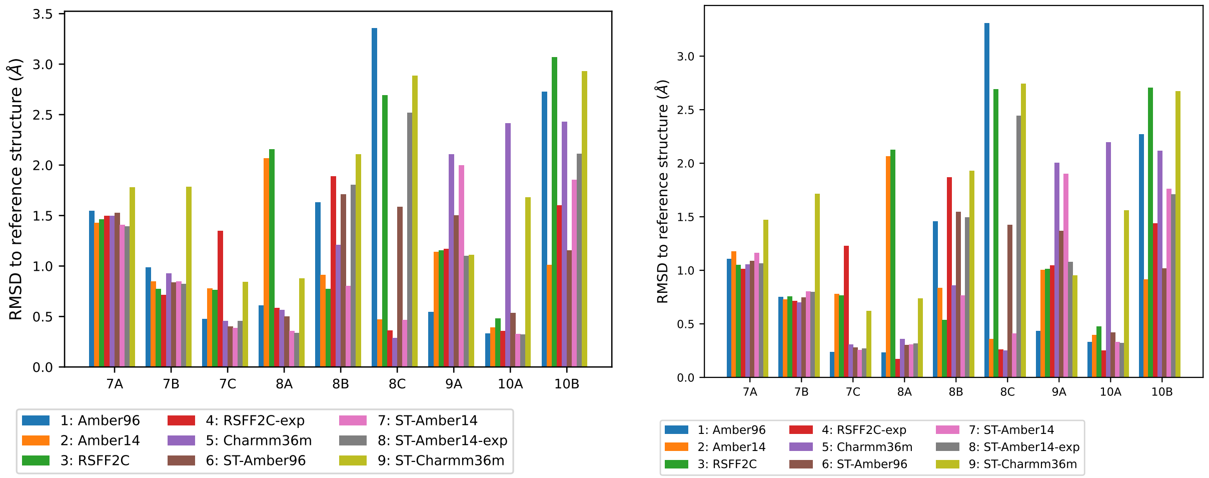

(a) (b)

Figure S5: (a) RMSD to the reference structure for the best ranked cluster for each peptide. Same figure as figure 10a of the article. (b) Minimum RMSD of the best ranked cluster to the 20 NMR models of each peptide.

### 6 Combination of four simulations for peptide 9A

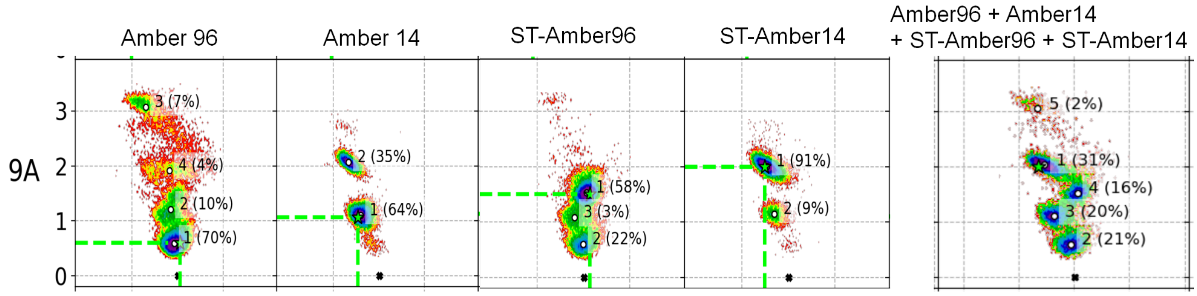

Figure S6: Example of combination of four simulations for peptide 9A.

### 7 Additional test set of four peptides

#### 7.1 Introduction

Table S1: Cyclic peptide test data set. All structures of the peptides in free form (i.e. not bound to a protein) have been solved by X-ray.[4] Upper case letters are L-amino acids and lower case letters are D-amino acids.

| PDB code | number of amino acids | sequence | atoms |
|---|---|---|---|
| 6UD9 | 8 | PkVepKvE | 134 |
| 6UCX | 8 | PeVkpEvK | 134 |
| 6UDZ | 10 | EppKvePPkV | 162 |
| 6UDW | 10 | qTRPDQtrpd | 162 |

We selected four additional peptides as test set. The peptides listed in table S1 are of the same nature as the nine initial peptides of the study, as they have the same number of amino acids, two of 8 A.A. and two of 10 A.A. and as they have a mixed chirality. All of them were designed peptides from the David Baker group,[4] but solved by X-ray instead of NMR. As can be seen from the sequences in table S1 and as described in the corresponding article,[4] the four peptides have an internal symmetry, more exactly an improper S2 rotational symmetry.

#### 7.2 Individual and combined free energy maps

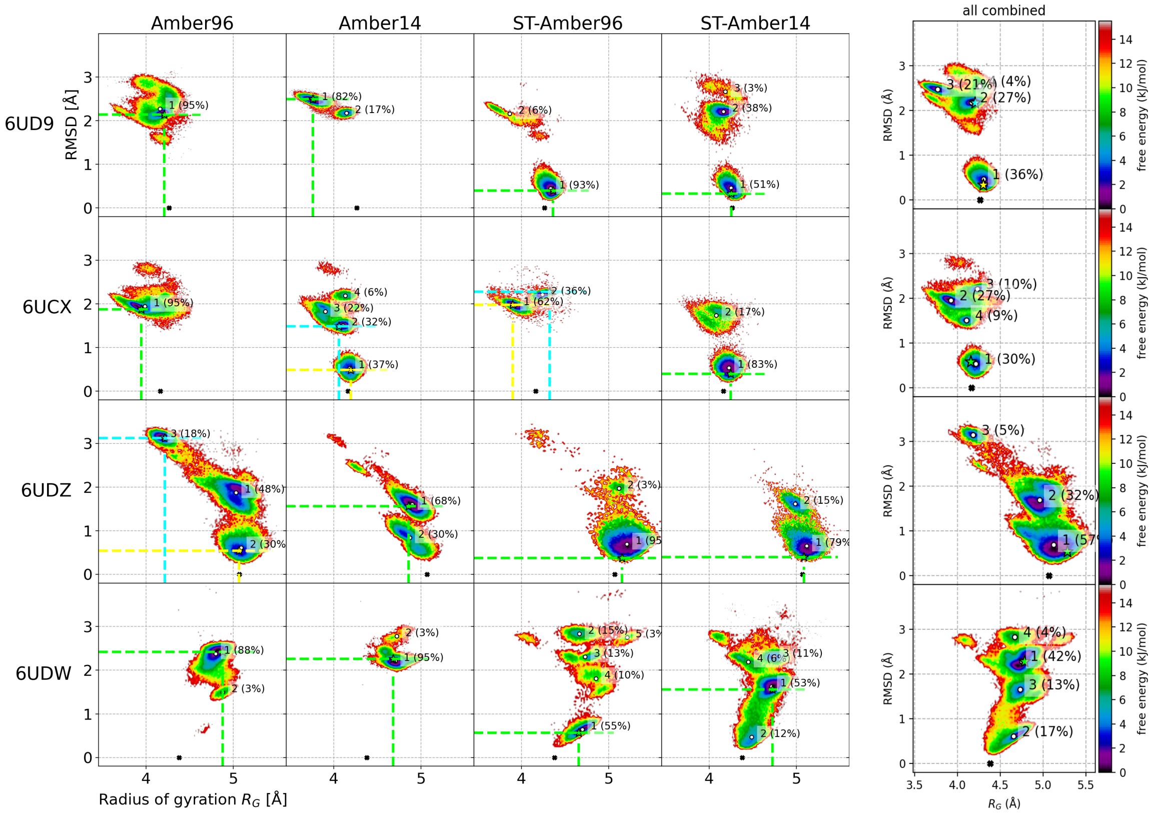

Figure S7: Free energy maps projected onto the radius of gyration (horizontal axis) and the RMSD to the reference structure. The units of the axes are in Å and the free energy unit is in kJ/mol. The first two columns were obtained with the REMD protocol and the next two columns with the ST protocol. The force field employed is indicated on top of each column. All columns are implicit solvent simulations. The last column combines all four simulations. The clustering results are indicated by the cluster ids and their fraction of the total number of frames, from the most populated cluster (id = 1) to the less populated cluster (clusters with a size below 2% are not shown). The centroid of each cluster is indicated by white dots. The centroid position of the most populated and lowest free energy cluster of the PCA projection (see Figure S9) is indicated by a green star. In the rare case that the position of the lowest free energy value in the PCA maps is different from the position of the most populated cluster, we plotted a cyan star for the former and a yellow star for the latter. The radius of gyration of the reference experimental structure is indicated by a black cross on the 0.0 RMSD line.

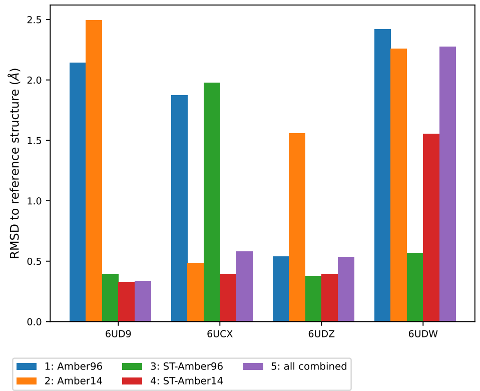

Figure S8: RMSD to the reference structure for the best ranked cluster for each peptide.

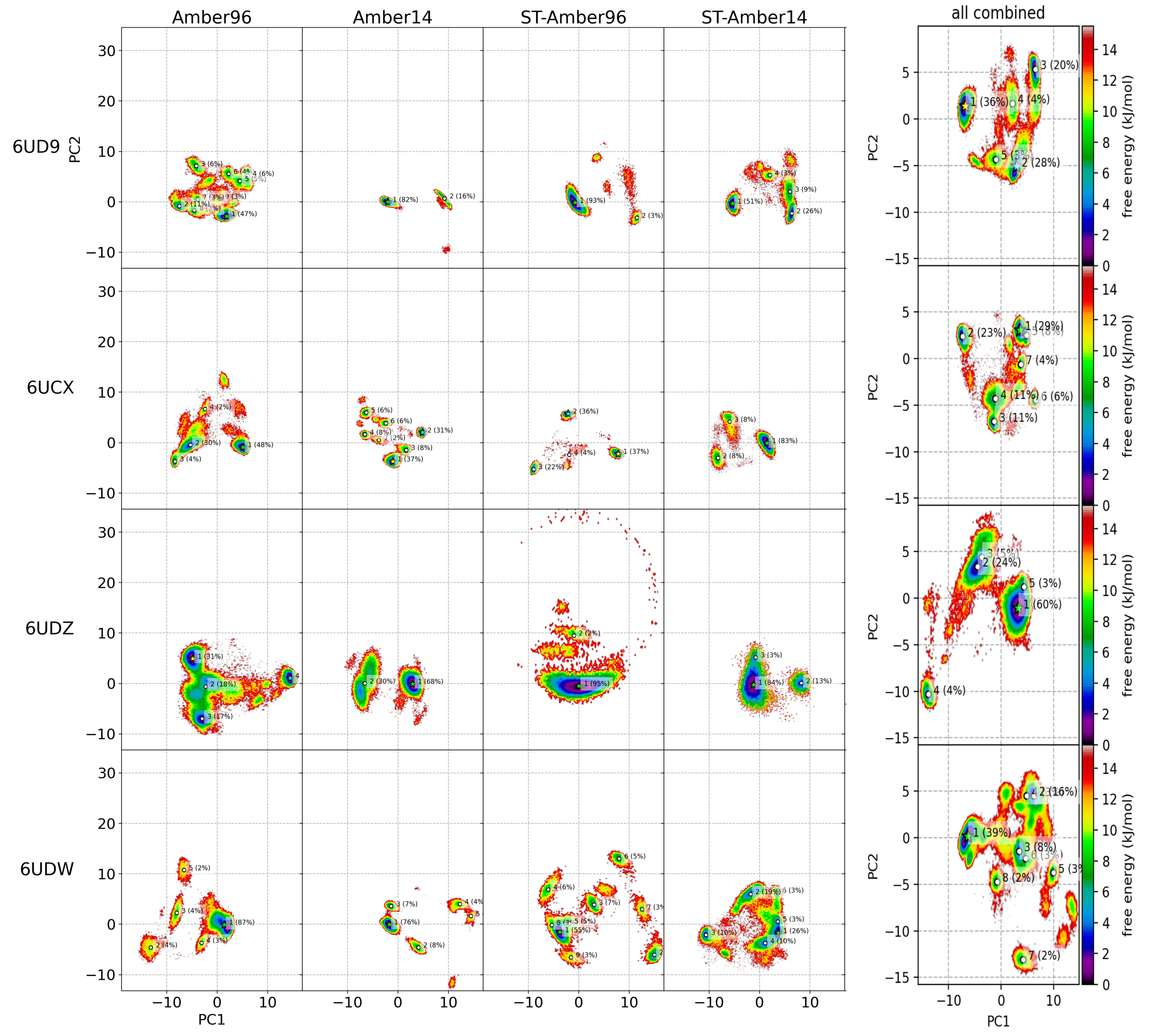

Figure S9: Free energy maps projected onto the first two principal components. The units of the axes are arbitrary and the free energy unit is in kJ/mol. The first two columns were obtained with the REMD protocol and the last two columns with the ST protocol. The force field employed is indicated on top of each column. All columns are implicit solvent simulations. The last column combines all four simulations. The clustering results are indicated by the cluster ids and their fraction of the total number of frames, from the most populated cluster (id = 1) to the less populated cluster (clusters with a size below 2% are not shown). The centroid of each cluster is indicated by white dots. The centroid position of the most populated and lowest free energy cluster is indicated by a green star. In the rare case that the position of the lowest free energy value is different from the position of the most populated cluster, we plotted a cyan star for the former and a yellow star for the latter.

### 8 Special case: Peptide 8A

#### 8.1 Introduction

Looking at the free energy maps of figure 9 of the article, one can see that the peptide 8A is a special case. The REMD Amber14 and RSSF2C simulations are the only simulations that sample an unique cluster at 2.2Å. None of the other simulations sample this cluster at all, which is not observed for the other peptides.

The current section presents the additional simulations and analysis we made to gain a deeper understanding to what has happened here for the peptide 8A. First we extracted the centroid structure of this unique cluster at 2.2Å, named here ”clust1”. We then performed REMD and ST simulations starting from the clust1 structure. We employed the Amber96, Amber14 and Charmm36m force fields, all with an implicit solvent. In addition we modified the Amber14 force field (”Amber14 mod”), which is explained below. For the REMD simulations, we produced 100 ns for each of the eight replica. For the ST simulations, 10 $\mu s$ have been produced.

The obtained free energy maps are shown in figure S10. The starting structure ”clust1” is indicated by an orange circle. As expected the REMD Amber14 simulation stays the whole 100 ns simulation very near to clust1. As before, the Amber96 and Charmm36m force fields do not stay near clust1. More surprisingly the ST-Amber14 simulation left the clust1 basin very quickly.

To understand better the reasons behind these observations, we plotted the backbone dihedrals $\phi$ (figure S11), $\psi$ (figure S10) and $\omega$ (figure S12). Only for the $\psi$ dihedral angles some significant differences are observed among the simulations. Especially the first two $\psi$ angles of the first two residues 1ASP and 2ASP show significant changes: for Amber14 the first one oscillates between $\pm$ 180 degrees, while for Amber96 it first do the same, but very quickly stabilizes at about -20 degrees. For ST-Amber14 this stabilization occurs even quicker, right from the first few ns. More importantly, the second $\psi$ angle remains constant in Amber14 at about -10 degrees, while it progressively moves to about 120 degrees in Amber96 and very quickly in ST-Amber14. The values of these first $\psi$ angles are 164 and -12 degrees for clust1, while they are -21 and 128 degrees for the experimental reference structure. This confirms that a joint swap of first two $\psi$ angles has to happen to bring the clust1 structure nearer to the reference structure. A third

$\psi$ angle has to change: the one of 4THR, as it has to go from -45 to 112 degrees from clust1 to the reference structure. This is effectively observed in the plots of the $\psi$ angles for Amber96 and again very quickly for ST-Amber14. The ST-Amber96 $\psi$ angle curves are very similar to the ones of ST-Amber14, the "jumps" happened here visibly already in the equilibration phase.

The Charmm36m REMD simulation behaves here similar to the Amber96 REMD simulation. The ST-Charmm36m results are quite different from all other simulations, as can be seen from the free-energy map and also from the $\phi$ and $\psi$ angle curves, which show a high flexibility on all residues, except 4THR.

To understand better where the differences between Amber96 and Amber14 are coming from, we replaced in Amber14 the $\psi$ angle force field parameters with the ones from Amber96. This allowed the "Amber14 mod" simulation to escape the clust1 basin with the same speed as with Amber96 (compare the 2ASP $\psi$ angles), but while keeping a preference for the clust1 basin and swapping between both. This produced a free energy map that is effectively a mixture of the ones of Amber96 and Amber14. This confirms that the energy restraints on the $\psi$ angles are the main driver on the observed differences, at least when comparing Amber96 with Amber14 in REMD.

We observed in the clust1 structure a salt bridge between 2ASP and 6DARG side chains and wondered if this salt bridge might stabilize or even constrain the clust1 structure. This salt-bridge is only conserved in the REMD Amber14 simulation, see figure S13. To check the importance of this salt bridge, we mutated *in-silico* the 6DARG to 6DALA and performed six REMD 100 ns simulations with Amber96, Amber14 and Charmm36m and with two runs. The free energy maps and the $\psi$ angles of the Amber14 simulation show that even without this salt bridge, the Amber14 simulations do not escape the clust1 basin, see figure S14. This confirms the importance of the $\psi$ energy terms, or generally backbone dihedral terms for cyclic peptides.

All this do not explain why the ST-Amber14 simulation escape so quickly the clust1 basin. To investigate this question, we tried to "boost" the REMD Amber14 simulation to be able to escape the clust1 basin quicker. Therefore we increased the maximum temperature to 500, 600 or 700K, while keeping the same number of eight replica. We performed a 100 ns REMD simulation on each of the three maximum temperatures in Amber14 and starting from clust1. Figure S15 shows that while 500K is not sufficient to escape the basin in only 100 ns, at 600K this happens after 20 ns and at 700K after 5 ns. It shows also that 100 ns are not sufficient to propagate the low RMSD conformations from the high temperature replica to the low temperature replica in a significant manner. Perhaps a higher number of replica would be necessary for this. The ST-Amber14 simulation has only a maximum temperature of 500K, but as ST is a completely different method compared to REMD, other factors may allow ST to escape the clust1 basin very quickly.

To conclude, the observation of a highly populated clust1 basin in REMD Amber14 and RSFF2C seems to be due to the $\psi$ force field parameters of Amber14 and by extension also RSFF2C which is derived from Amber14. These differences are also observed in conventional MD simulations at 300K, see figure S16. The differences between REMD and ST on Amber14 are mainly due to the different capacity to escape this energy basin more or less quickly.

#### 8.2 Backbone dihedrals

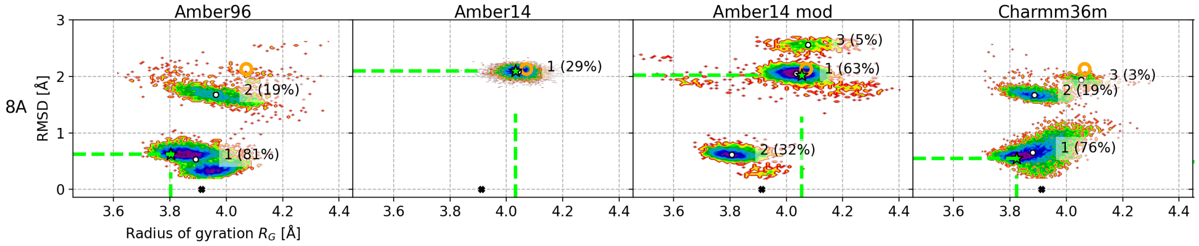

(a) REMD

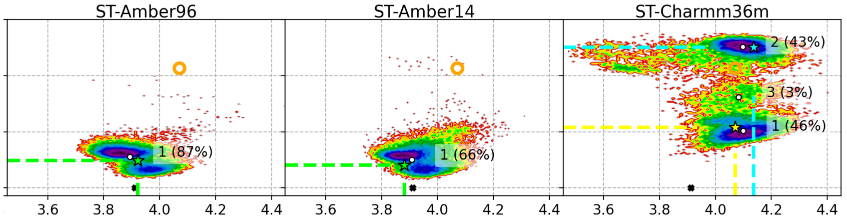

(b) ST

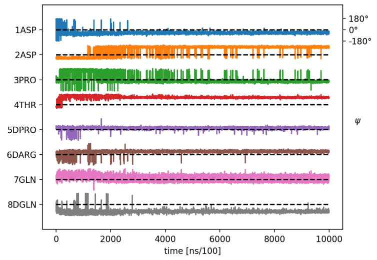

(c) Amber96

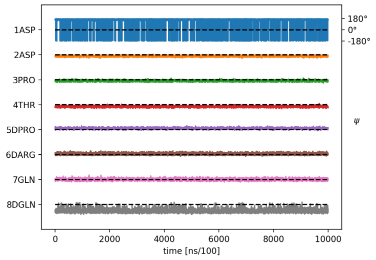

(d) Amber14

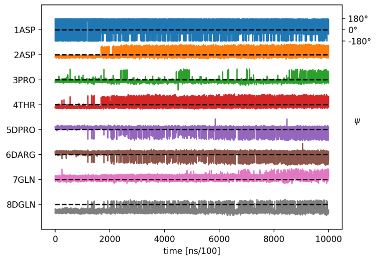

(e) Amber14 mod

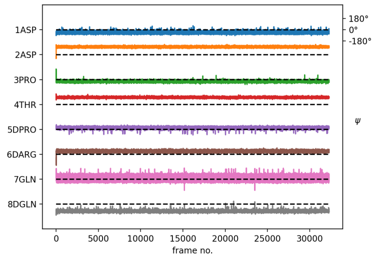

(f) ST-Amber96

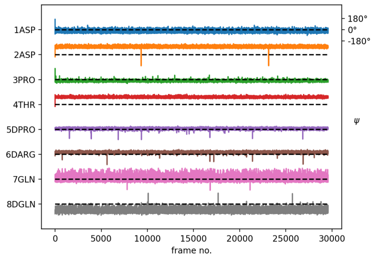

(g) ST-Amber14

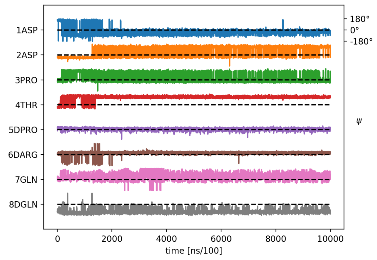

(i) Charmm36m

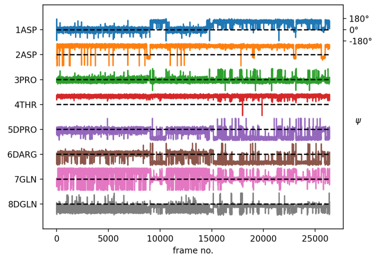

(j) ST-Charmm36m

Figure S10: Special case peptide 8A. All simulations started from the first cluster at 2.2Å from the reference structure only observed in Amber14 and RSFF2C. (a)-(b) Free energy maps. The starting structure is indicated by an orange circle. (c)-(j) $\psi$ backbone dihedrals.

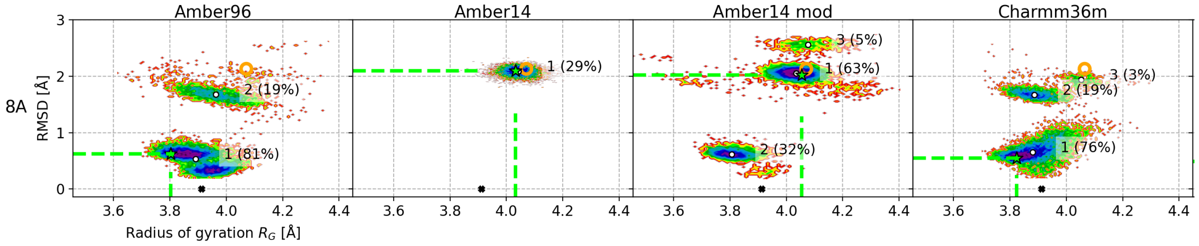

(a) REMD

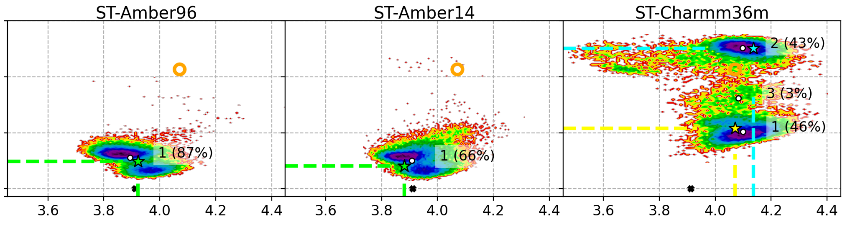

(b) ST

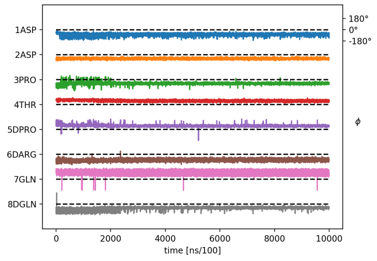

(c) Amber96

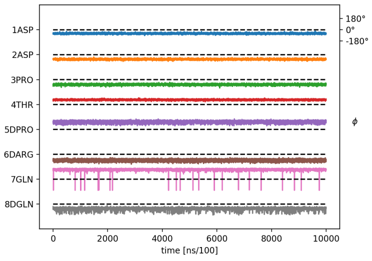

(d) Amber14

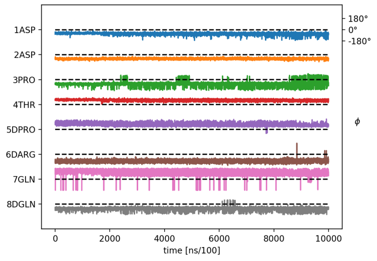

(e) Amber14 mod

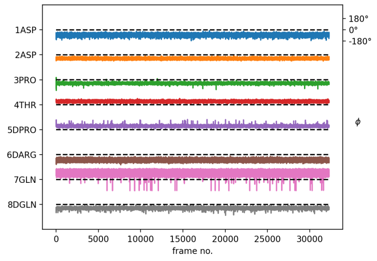

(f) ST-Amber96

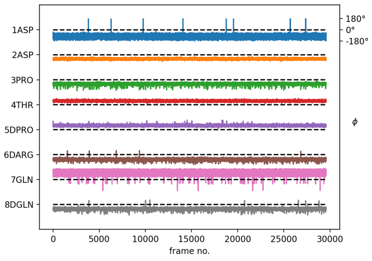

(g) ST-Amber14

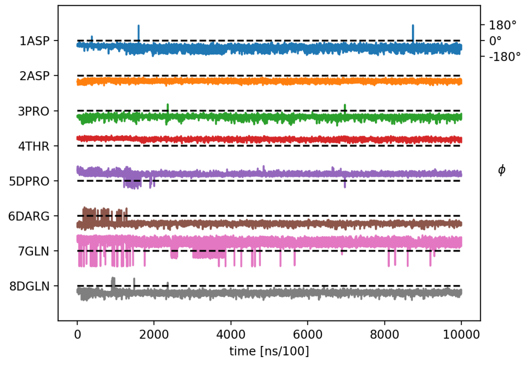

(i) Charmm36m

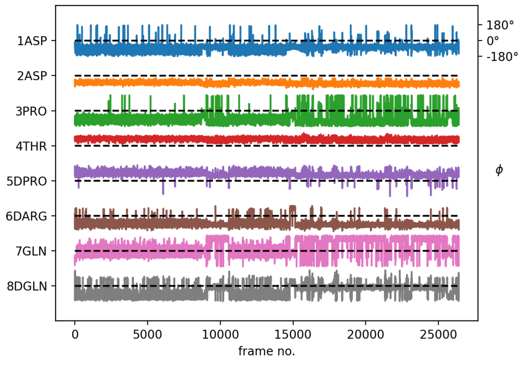

(j) ST-Charmm36m

Figure S11: Special case peptide 8A. All simulations started from the first cluster at 2.0Å from the reference structure only observed in Amber14 and RSFF2C. (a)-(b) Free energy maps. The starting structure is indicated by an orange circle. (c)-(j) $\phi$ backbone dihedrals.

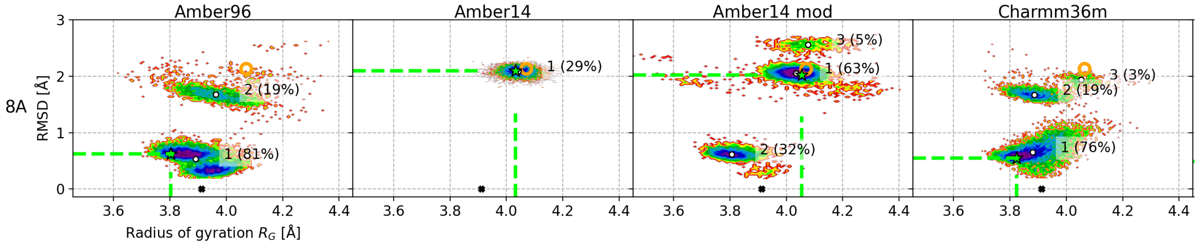

(a) REMD

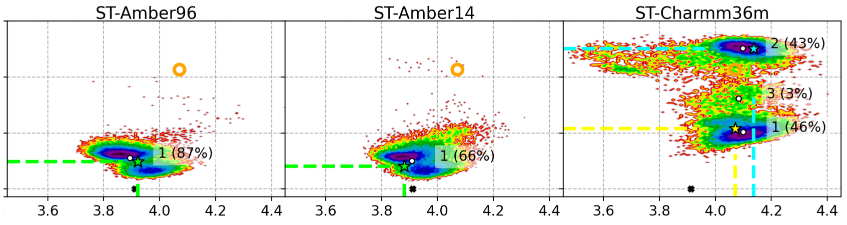

(b) ST

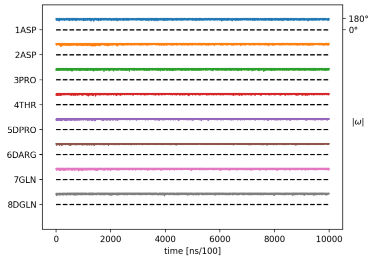

(c) Amber96

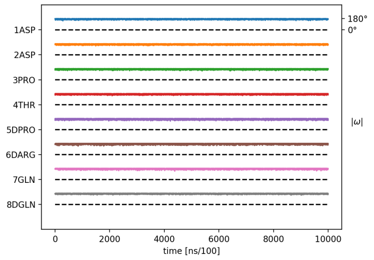

(d) Amber14

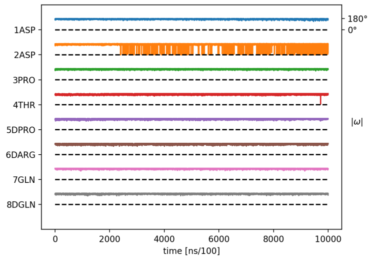

(e) Amber14 mod

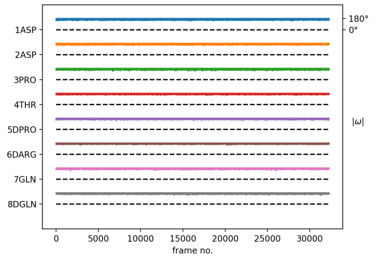

(f) ST-Amber96

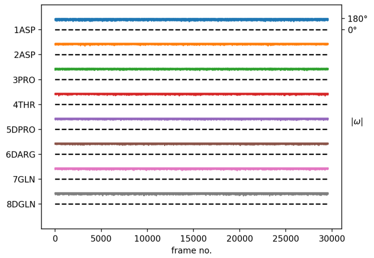

(g) ST-Amber14

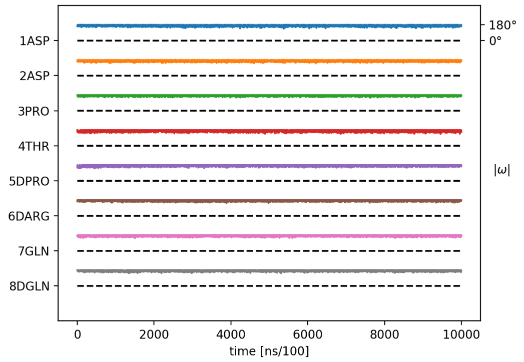

(i) Charmm36m

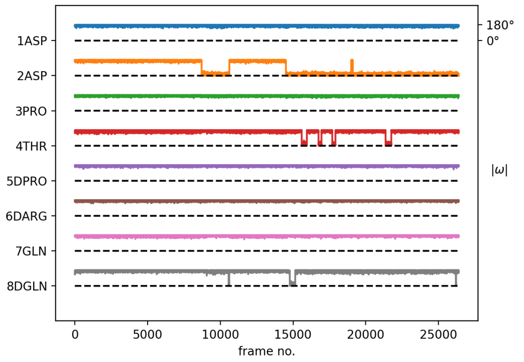

(j) ST-Charmm36m

Figure S12: Special case peptide 8A. All simulations started from the first cluster at 2.0Å from the reference structure only observed in Amber14 and RSFF2C. (a)-(b) Free energy maps. The starting structure is indicated by an orange circle. (c)-(j) $\omega$ backbone dihedrals.

#### 8.3 Salt bridge 2ASP - 6DARG with Amber14

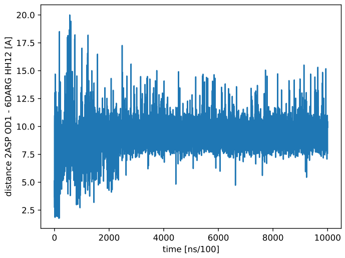

(a) Amber96

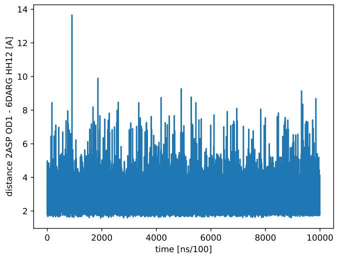

(b) Amber14

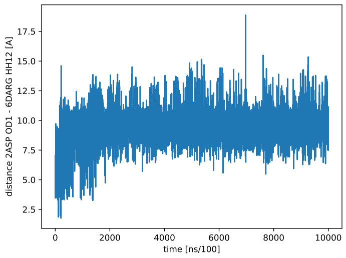

(c) Charmm36m

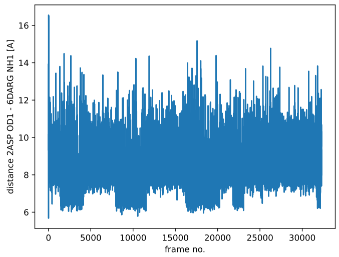

(d) ST-Amber96

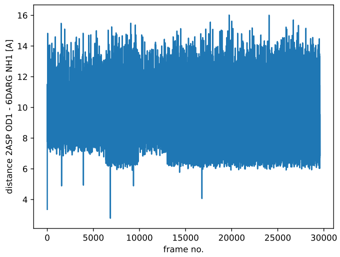

(e) ST-Amber14

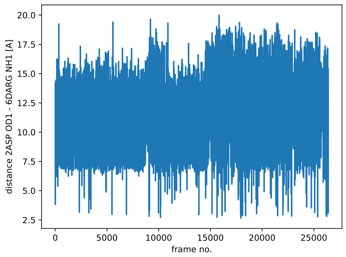

(f) ST-Charmm36m

Figure S13: Special case peptide 8A. With Amber14 in REMD a stable salt bridge is observed in the first cluster "clust1" between the side-chains of 2ASP and 6DARG, but not in the other force fields, nor with ST.

#### 8.4 Mutant 6DARG to 6DALA

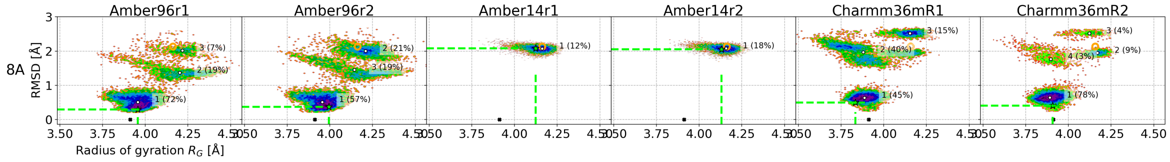

(a)

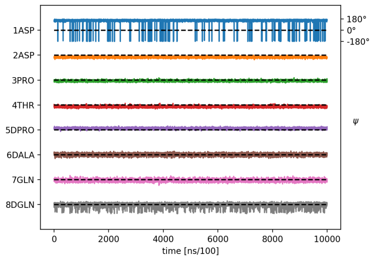

(b)

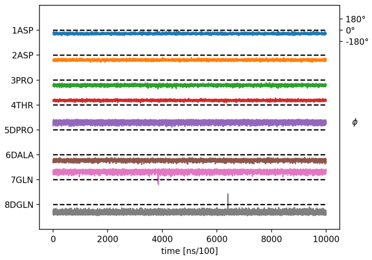

(c)

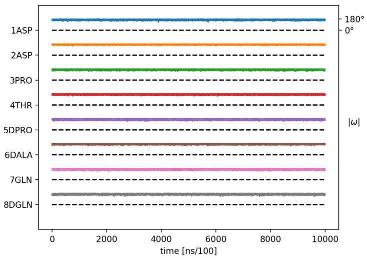

(d)

Figure S14: Special case peptide 8A, mutated at 6DARG to 6DALA in order to remove the salt bridge between 2ASP and 6DARG observed in the first cluster "clust1". (a) Free energy maps of REMD simulations with two runs per force field (r1 and r2). (b)-(d) Backbone dihedrals of the Amber14 REMD simulation.

#### 8.5 Amber14: Cross barrier with higher temperatures in REMD

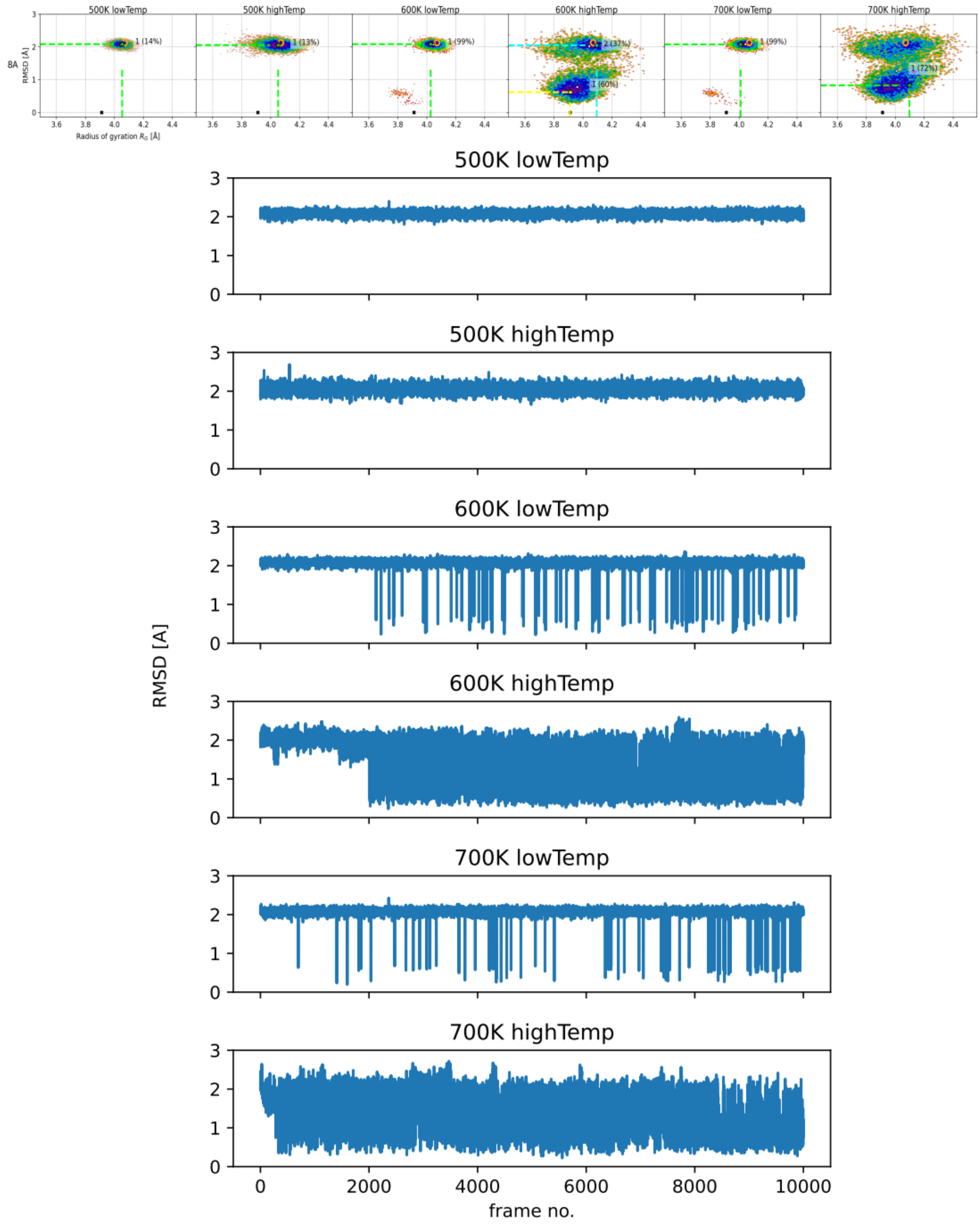

Figure S15: Special case peptide 8A with Amber14. All simulations started from the first cluster at 2.0Å from the reference structure only observed in Amber14 and RSFF2C. To cross the barrier with Amber14 the maximum temperature in REMD is increased to 500, 600 or 700K. The minimum temperature is in all cases 300K. The starting structure is indicated by an orange circle in the free energy maps. The maximum temperature of REMD is indicated in the titles. "LowTemp" = 300K, "HighTemp" = 500, 600 or 700K.

#### 8.6 Amber96 vs Amber14 in conventional MD at 300K

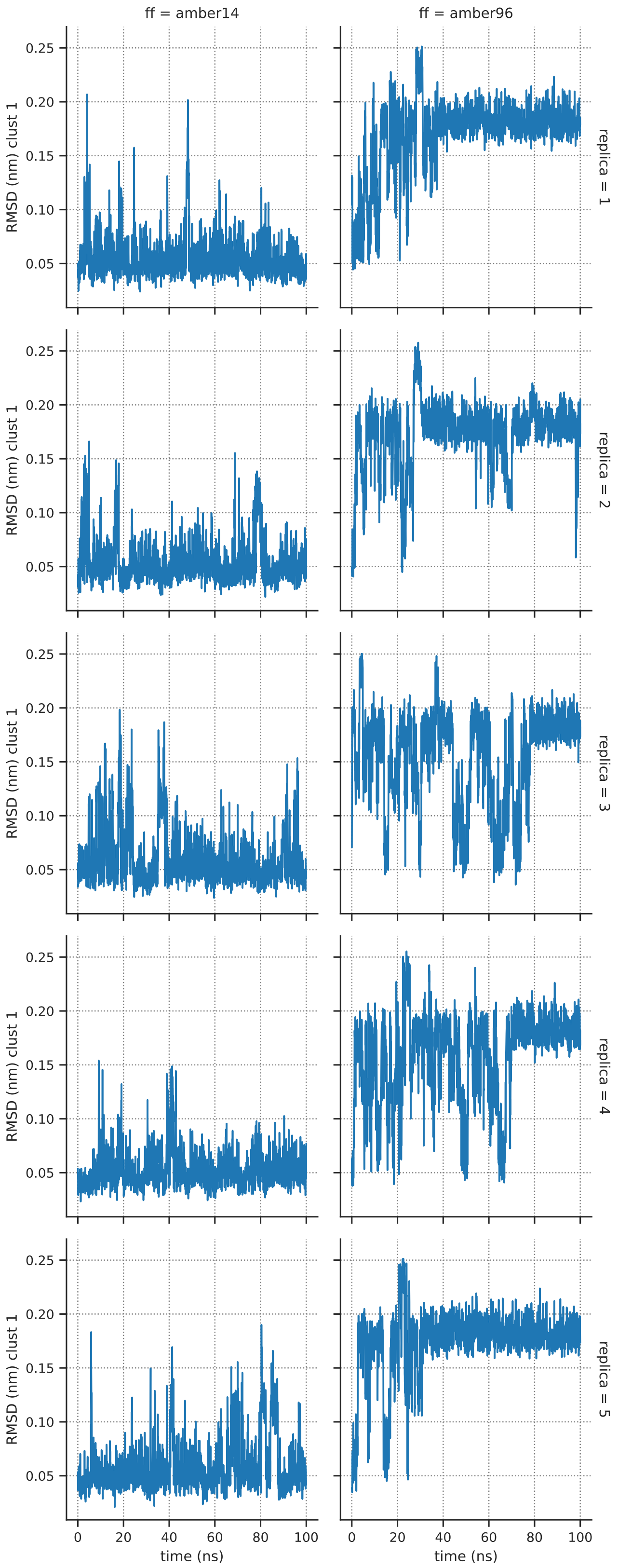

Figure S16: Special case peptide 8A with Amber96 and Amber14 in conventional MD at 300K repeated five times. All simulations started from the first cluster (named "clust1") at 2.0Å from the reference structure only observed in Amber14 and RSFF2C. The RMSD to this starting structure is plotted here.

### 9 Combined simulations

#### 9.1 All nine simulations

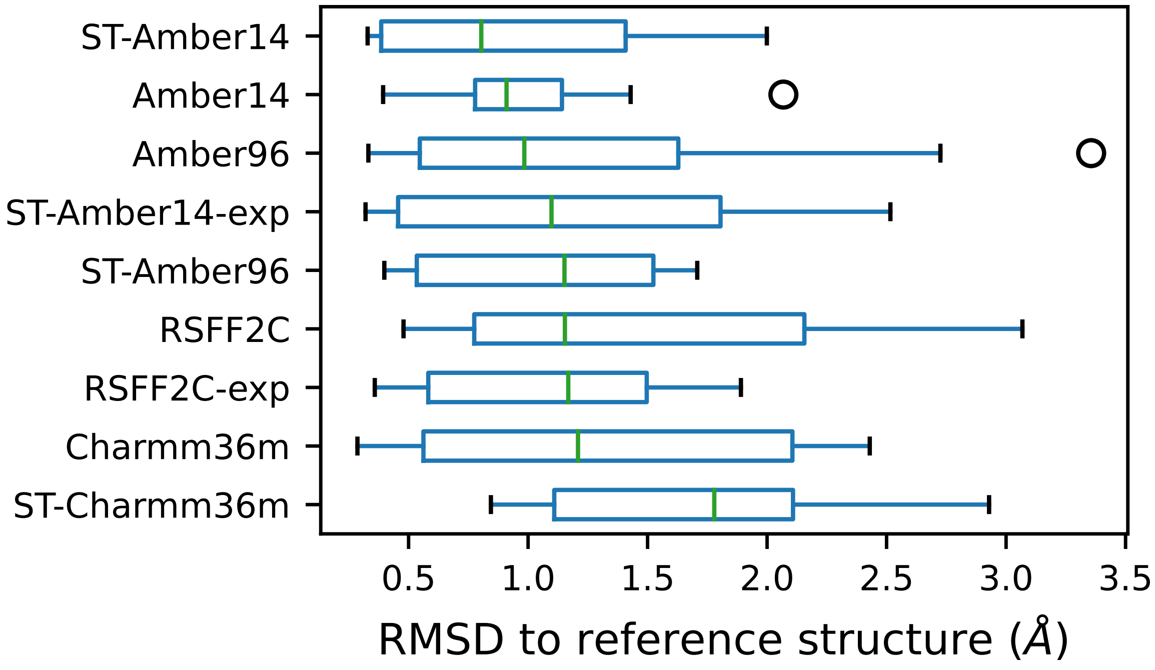

Figure S17: The nine single simulations used here for the combinations. RMSD to the reference structure for the best ranked cluster for each simulation. The RMSD values of the nine peptides are assembled into a box plot. The simulations are sorted here with increasing median RMSD values, shown with green lines.

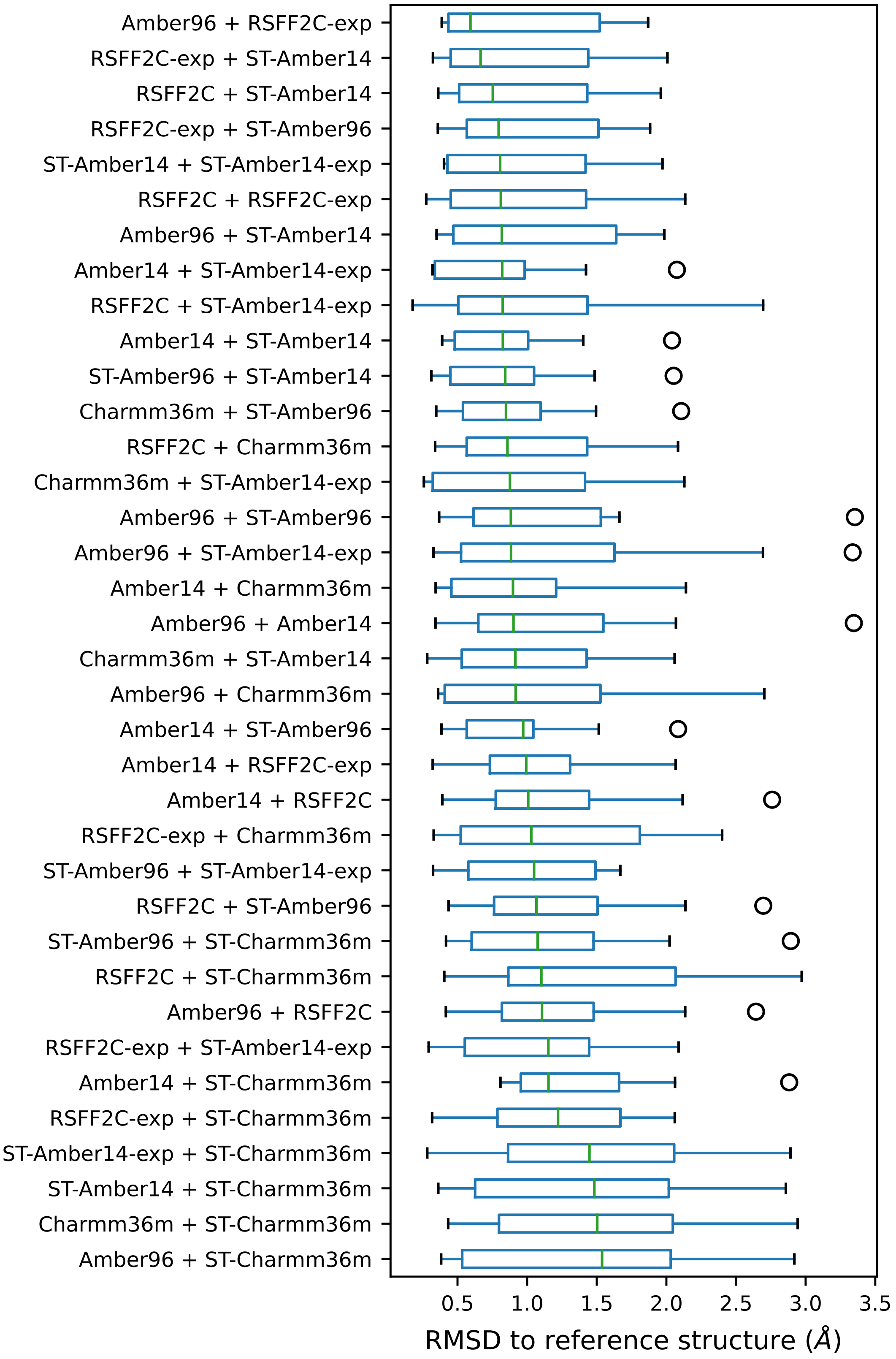

Figure S18: Two simulations combined. RMSD to the reference structure for the best ranked cluster for each combination. For each combination of simulations the RMSD values of the nine peptides are assembled into a box plot. The simulations are sorted here with increasing median RMSD values, shown with green lines.

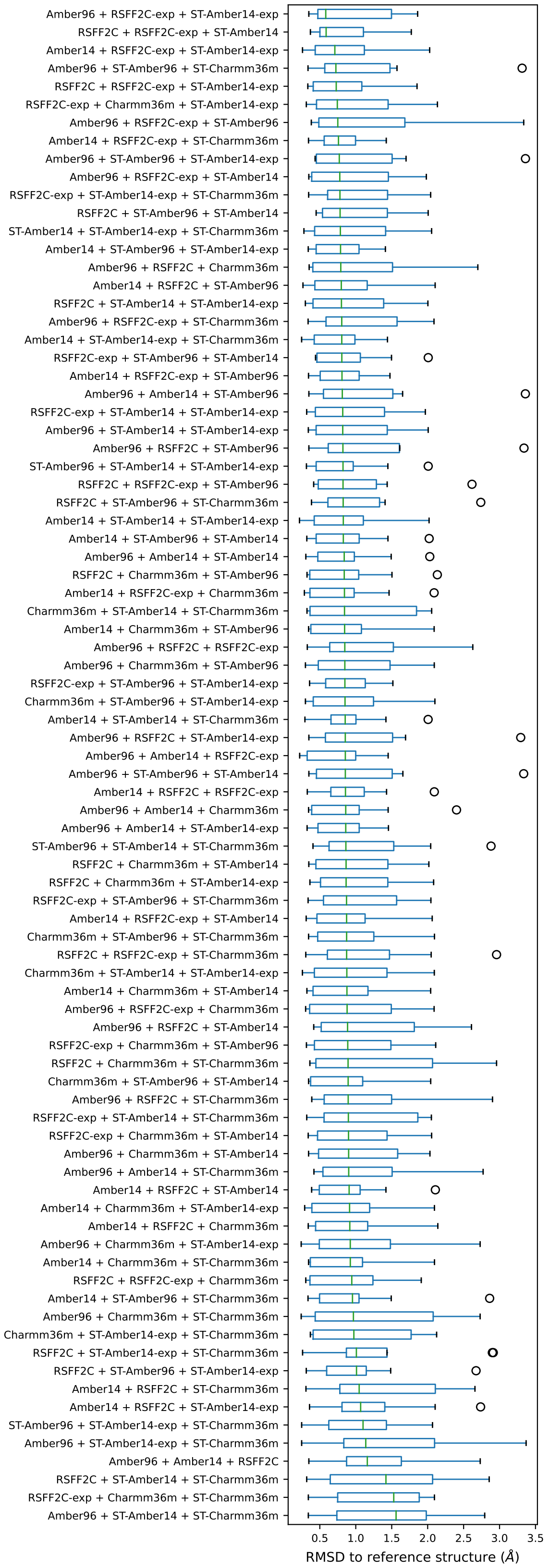

Figure S19: Three simulations combined. RMSD to the reference structure for the best ranked cluster for each combination. For each combination of simulations the RMSD values of the nine peptides are assembled into a box plot. The simulations are sorted here with increasing median RMSD values, shown with green lines.

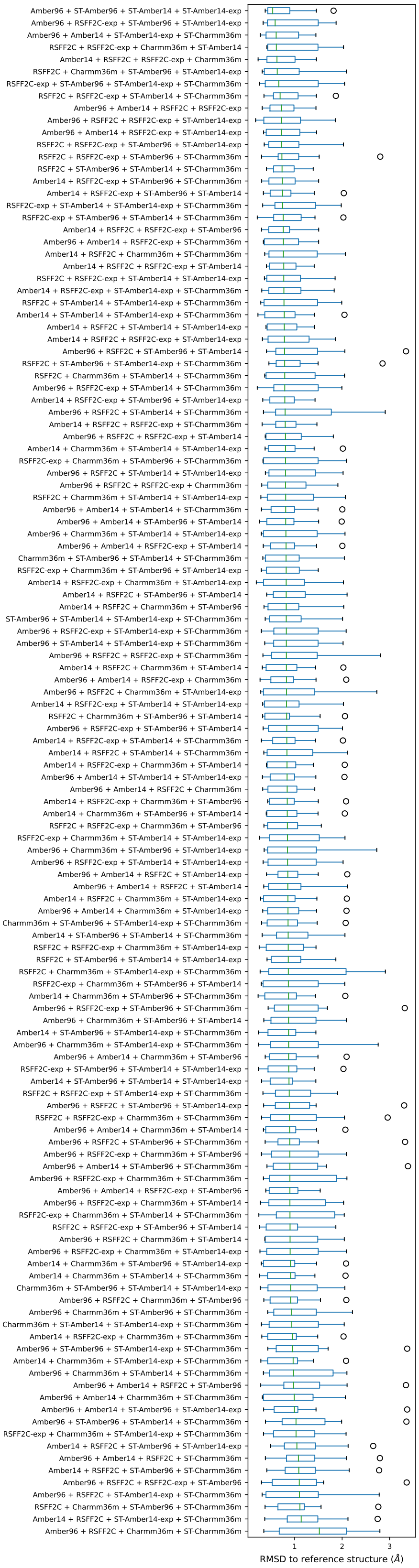

Figure S20: Four simulations combined. RMSD to the reference structure for the best ranked cluster for each combination. For each combination of simulations the RMSD values of the nine peptides are assembled into a box plot. The simulations are sorted here with increasing median RMSD values, shown with green lines.

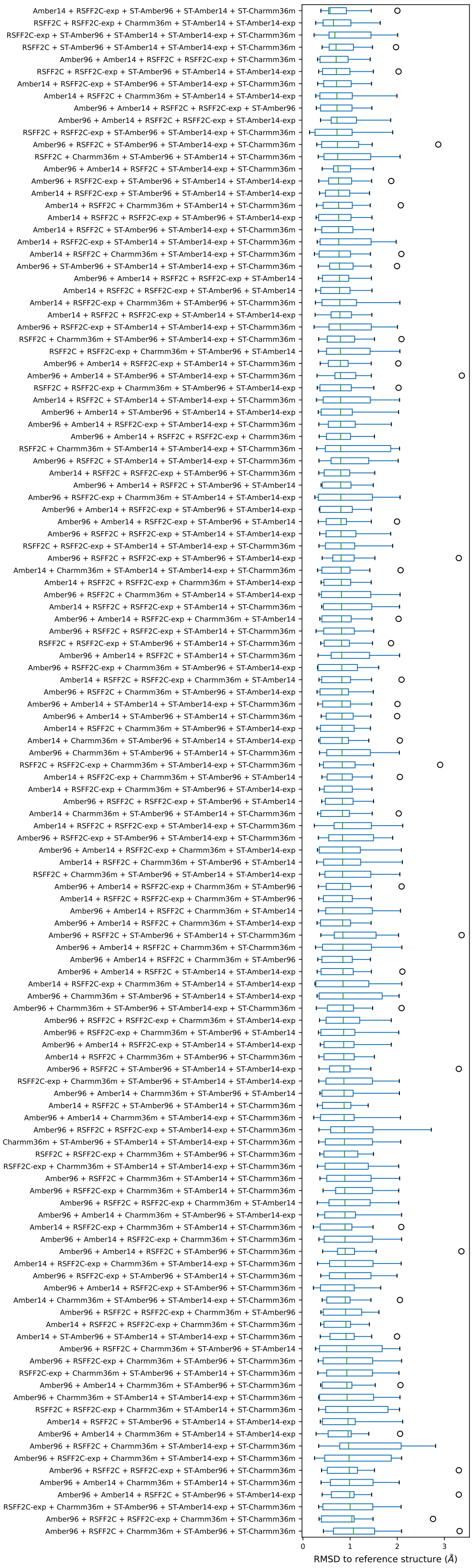

Figure S21: Five simulations combined. RMSD to the reference structure for the best ranked cluster for each combination. For each combination of simulations the RMSD values of the nine peptides are assembled into a box plot. The simulations are sorted here with increasing median RMSD values, shown with green lines.

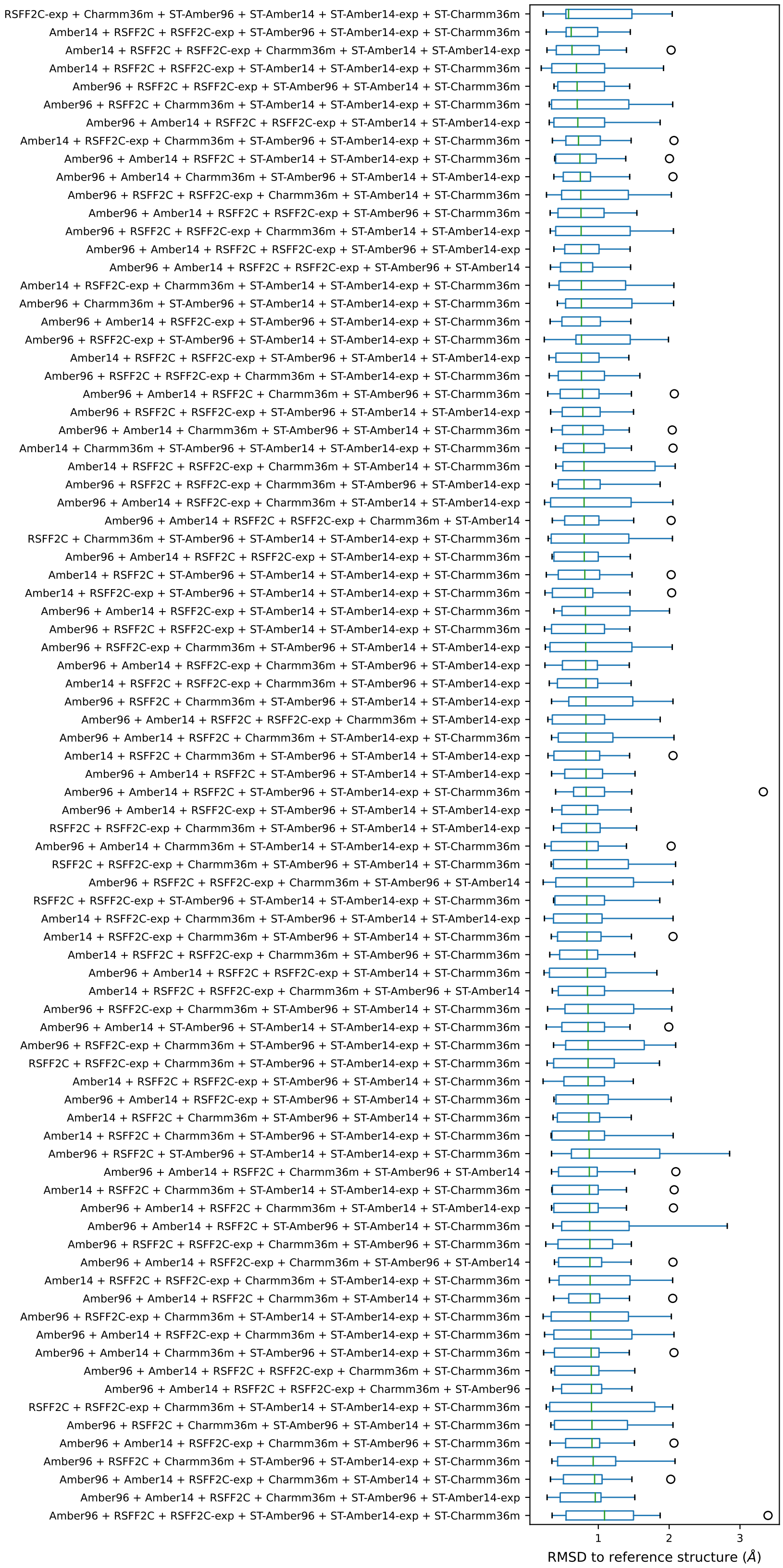

Figure S22: Six simulations combined. RMSD to the reference structure for the best ranked cluster for each combination. For each combination of simulations the RMSD values of the nine peptides are assembled into a box plot. The simulations are sorted here with increasing median RMSD values, shown with green lines.

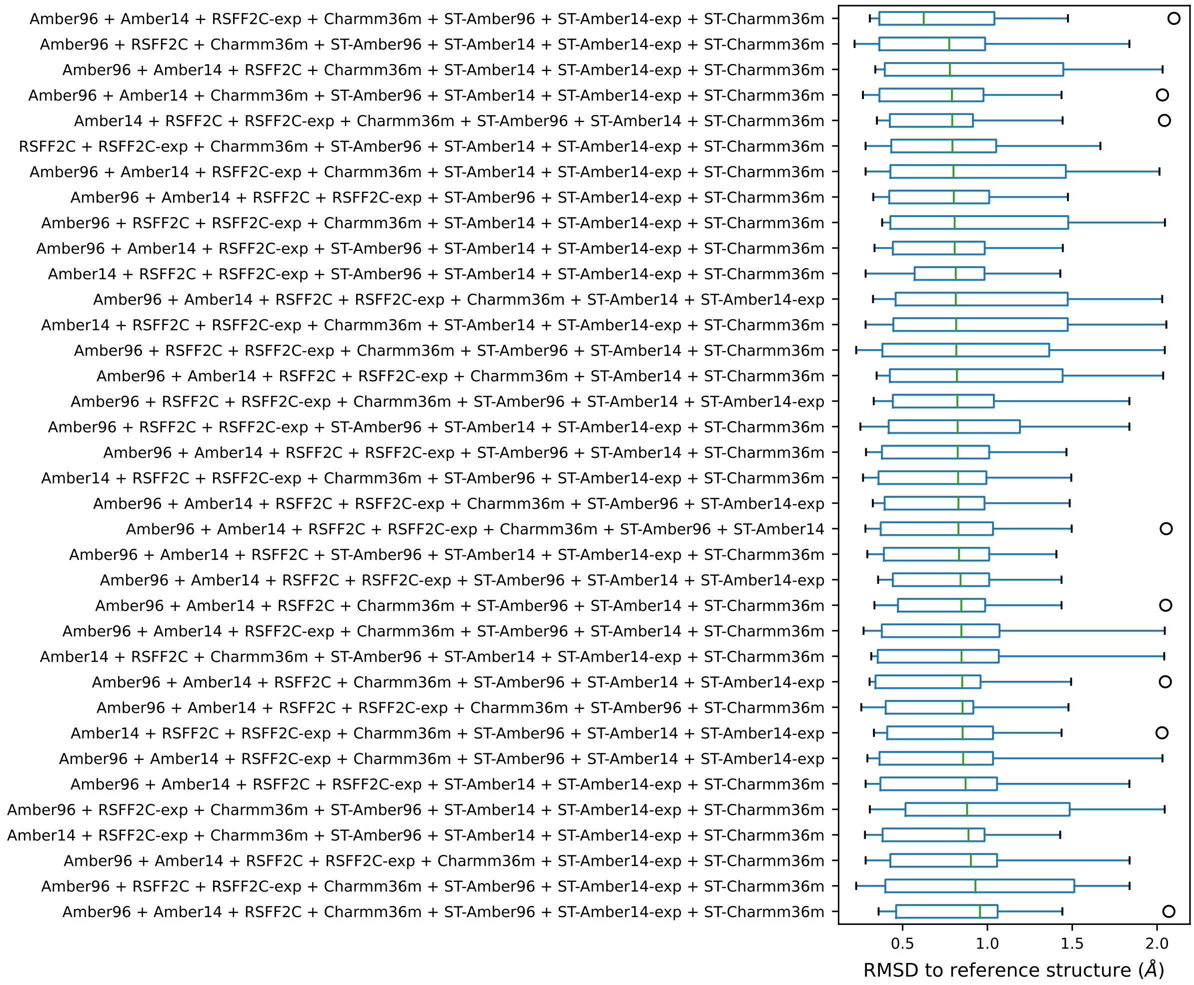

Figure S23: Seven simulations combined. RMSD to the reference structure for the best ranked cluster for each combination. For each combination of simulations the RMSD values of the nine peptides are assembled into a box plot. The simulations are sorted here with increasing median RMSD values, shown with green lines.

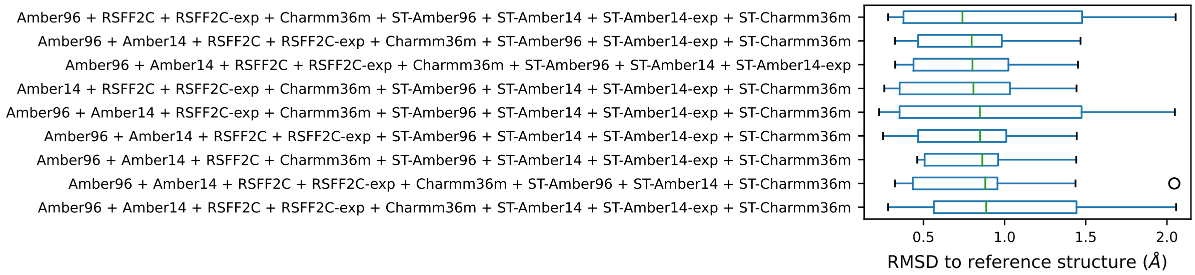

Figure S24: Eight simulations combined. RMSD to the reference structure for the best ranked cluster for each combination. For each combination of simulations the RMSD values of the nine peptides are assembled into a box plot. The simulations are sorted here with increasing median RMSD values, shown with green lines.

#### 9.2 Only implicit solvent simulations

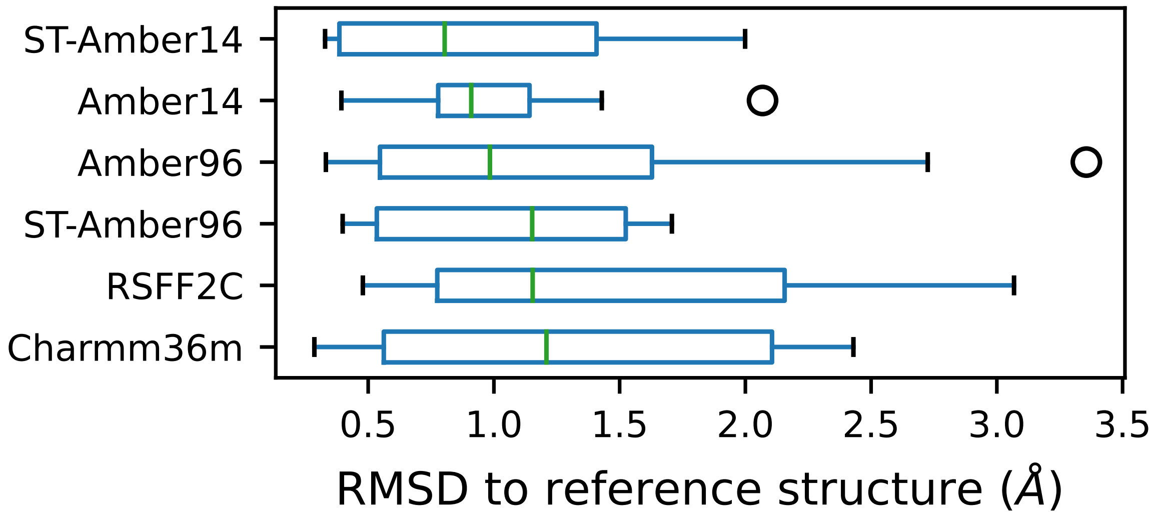

Figure S25: The six single implicit solvent simulations used here for the combinations. RMSD to the reference structure for the best ranked cluster for each simulation. The RMSD values of the nine peptides are assembled into a box plot. The simulations are sorted here with increasing median RMSD values, shown with green lines.

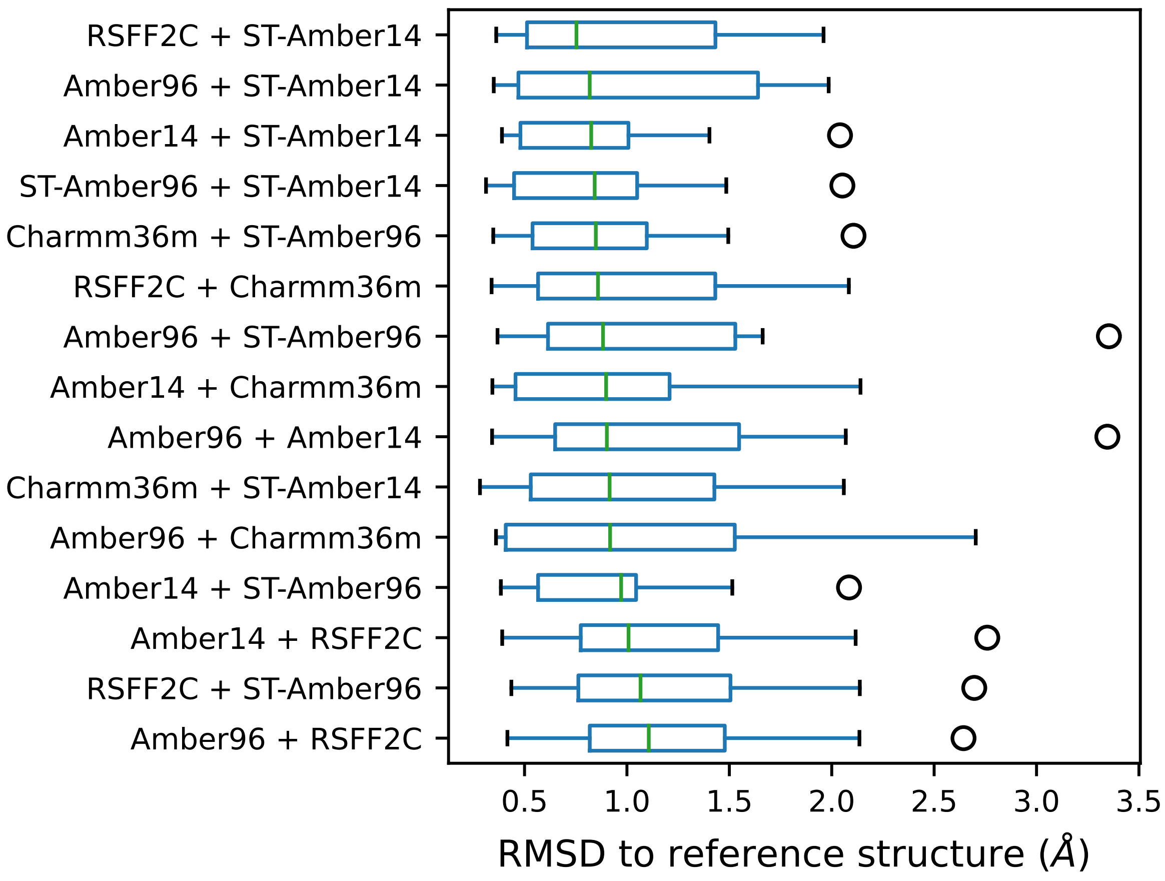

Figure S26: Two simulations combined. RMSD to the reference structure for the best ranked cluster for each combination. For each combination of simulations the RMSD values of the nine peptides are assembled into a box plot. The simulations are sorted here with increasing median RMSD values, shown with green lines.

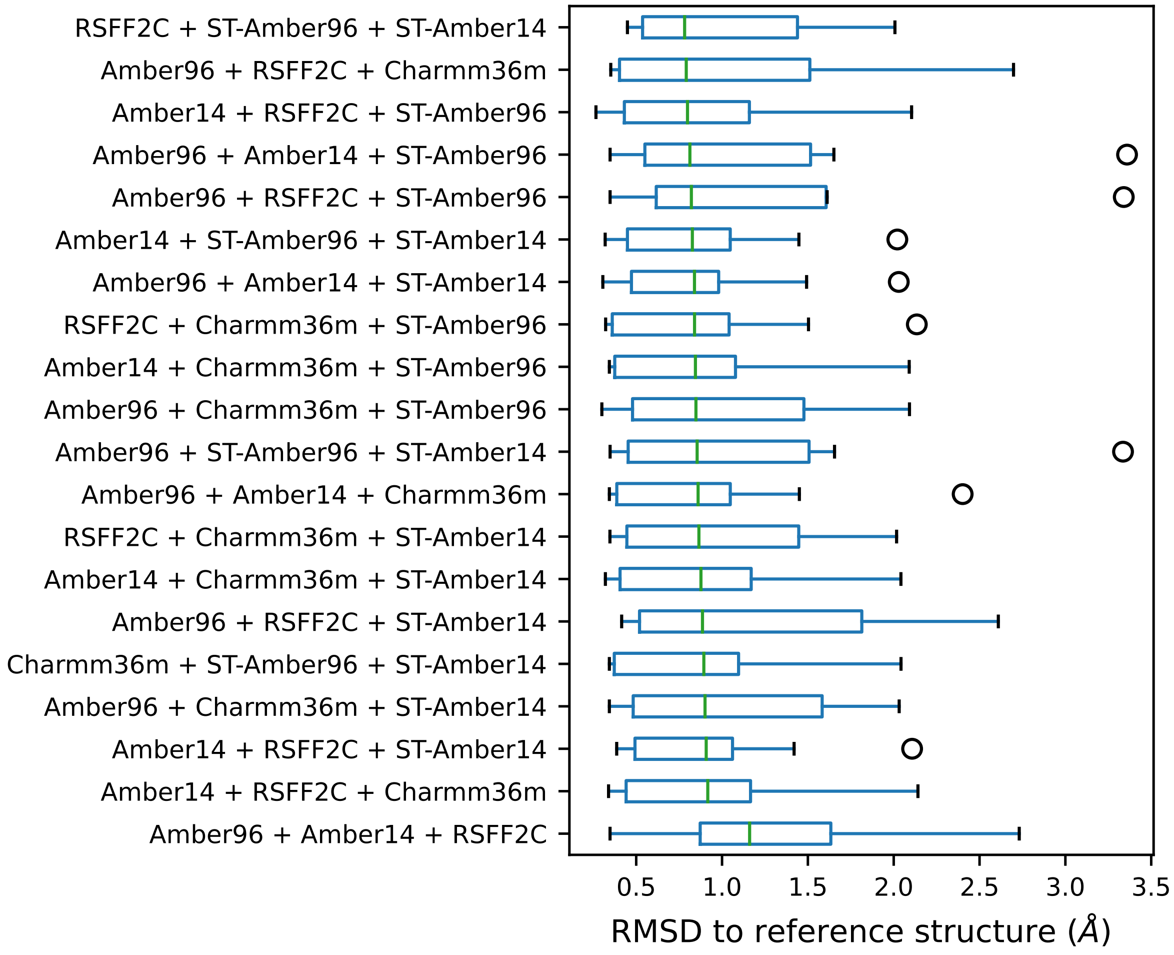

Figure S27: Three simulations combined. RMSD to the reference structure for the best ranked cluster for each combination. For each combination of simulations the RMSD values of the nine peptides are assembled into a box plot. The simulations are sorted here with increasing median RMSD values, shown with green lines.

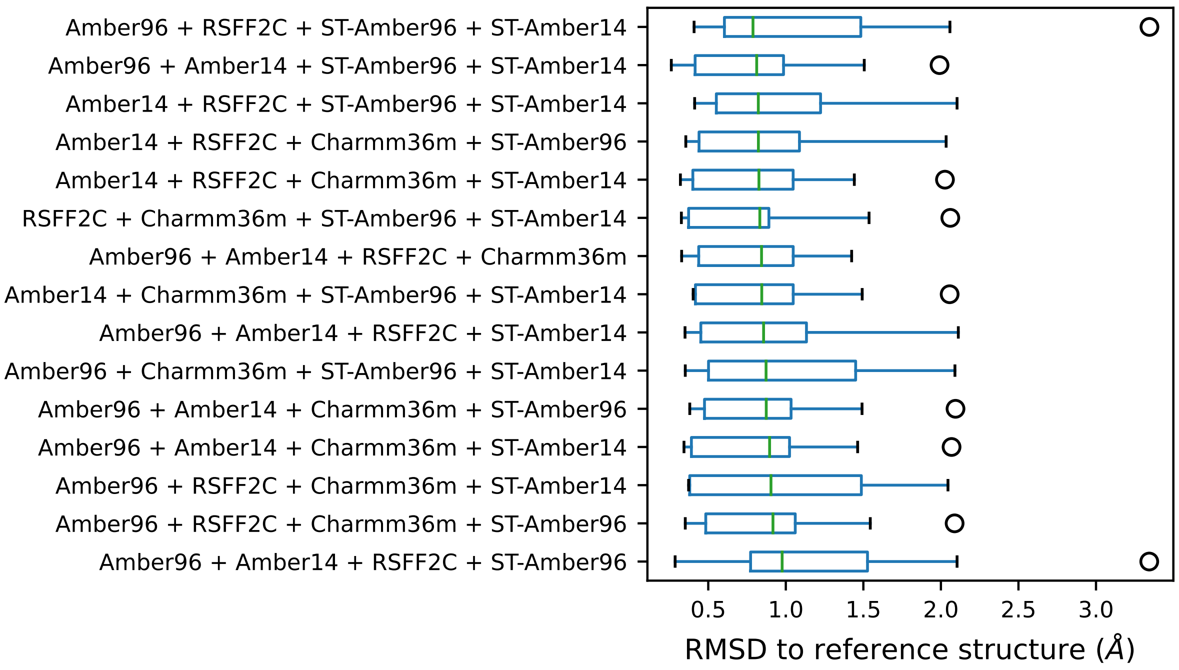

Figure S28: Four simulations combined. RMSD to the reference structure for the best ranked cluster for each combination. For each combination of simulations the RMSD values of the nine peptides are assembled into a box plot. The simulations are sorted here with increasing median RMSD values, shown with green lines.

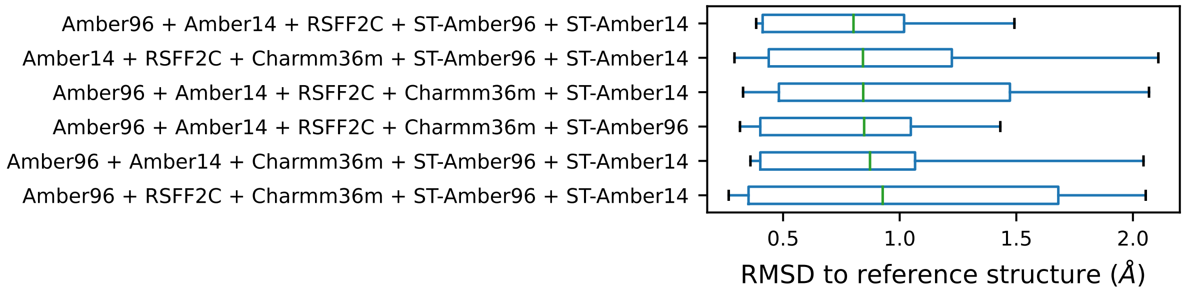

Figure S29: Five simulations combined. RMSD to the reference structure for the best ranked cluster for each combination. For each combination of simulations the RMSD values of the nine peptides are assembled into a box plot. The simulations are sorted here with increasing median RMSD values, shown with green lines.

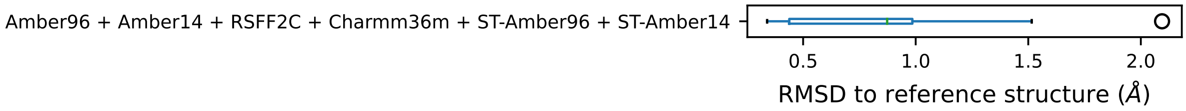

Figure S30: Six simulations combined. RMSD to the reference structure for the best ranked cluster for each combination. For each combination of simulations the RMSD values of the nine peptides are assembled into a box plot. The simulations are sorted here with increasing median RMSD values, shown with green lines.
